# Upstream of N-Ras C-terminal cold shock domains mediate poly(A) specificity in a novel RNA recognition mode and bind poly(A) binding protein during translation regulation

**DOI:** 10.1101/2022.11.26.518022

**Authors:** Nele Merret Hollmann, Pravin Kumar Ankush Jagtap, Johanna-Barbara Linse, Philip Ullmann, Marco Payr, Brice Murciano, Bernd Simon, Jochen S. Hub, Janosch Hennig

**Affiliations:** Structural and Computational Biology Unit, EMBL Heidelberg, Meyerhofstraße 1, 69117, Heidelberg, Germany; Collaboration for joint PhD degree between EMBL and Heidelberg University, Faculty of Biosciences; Chair of Biochemistry IV, Biophysical Chemistry, University of Bayreuth, Universitätsstrasse 30, 95447 Bayreuth, Germany; Theoretical Physics, Saarland University, 66123 Saarbrücken, Germany; Center for Biophysics, Saarland University, 66123 Saarbrücken, Germany

## Abstract

RNA binding proteins (RBPs) often engage multiple RNA binding domains (RBDs) to increase target specificity and affinity. However, the complexity of target recognition of multiple RBDs remains largely unexplored. Here we use Upstream of N-Ras (Unr), a multidomain RBP, to demonstrate how multiple RBDs orchestrate target specificity. A crystal structure of the three C-terminal RNA binding cold-shock domains (CSD) of Unr bound to a poly(A) sequence exemplifies how recognition goes beyond the classical π-π-stacking in CSDs. Further structural studies reveal several interaction surfaces between the N-terminal and C-terminal part of Unr with the poly(A)-binding protein (pAbp). This provides first atomistic details towards understanding regulation of translation initiation that is mediated by the interplay of these two proteins with each other and RNA.

## INTRODUCTION

RNA binding proteins (RBPs) interact with coding and non-coding RNAs as constitutive partners in ribonucleoprotein (RNP) complexes. The structural and mechanistic knowledge about RNP assemblies is scarce and mainly limited to large molecular machines, like the ribosome(1, 2), RNA polymerases(3–5) or the spliceosome(6–8). These machines are often highly abundant in cells and their target interaction is strong and constitutive, which is advantageous for mechanistic studies. However, many RBPs function as regulatory units, requiring transient and versatile interactions with their binding partners along with fluctuating abundance(9). Thereby these RBPs can respond to environmental changes or developmental cues quickly. The dynamic nature of RBPs is an advantage in their involvement in many regulatory pathways of the cell, including gene expression at all levels ranging from transcription, splicing, polyadenylation, localization, stabilization, and degradation to protein synthesis via their diverse roles in translation(10–13). This rather transient binding nature makes structural studies difficult and explains why RNA binding properties of most RBPs remain unexplored(12).

To ensure specific regulation through RBPs in the many different cellular processes, a certain RNA target specificity of the protein is a prerequisite. RBPs employ a set of RNA binding domains (RBDs) to engage their target RNAs. RBDs are often small and very conserved domains, with specificities towards single stranded (ssRNA) or double stranded RNA (dsRNA)(14) Although RBDs are the main drivers of protein-RNA interactions, the single domains are often not enough to discriminate target from non-target RNAs within the complex transcriptome of the cell(11). Most classical RBDs are around 10 kDa of size and can accommodate three to five contiguous RNA bases specifically(15), which is often not enough to endow RNA target recognition(11). Thus, a composition of multiple RBDs within one protein often increases specificity(16, 17). The majority of RBPs is composed of multiple RBDs, either of the same or of different domain types(11). This results in a large combinatorial variety of different domain classes and the diversity of architectures would influence the binding mode to the specific target RNA sequence.

One prerequisite to understand mechanistic details of the different binding modes of RBPs that induce target specificity, is the determination of RNP structures at an atomistic level. Over the years, there have been several efforts to examine structural features that dictate RNP binding specificity(18, 19). The knowledge of how single RBDs engage their target sequences increased and in some cases these studies offered insights into the role of multidomain arrangements in the recognition process(20, 21). However, the interplay of multiple RBDs in a single RBP is far from being understood and may change on a case-to-case basis.

Here we use *Drosophila* Unr, a multidomain RBP, to demonstrate the complexity of RNA binding. Unr is a highly conserved protein among metazoan, containing five ssRNA binding canonical cold-shock domains (CSD) and four additional non-canonical CSDs (ncCSDs), which lack the conserved RNA binding residues and are therefore incapable of independent RNA binding(22). More than hundreds of transcripts could be identified in previous co-immunoprecipitation studies with Unr(22, 23), reflecting its widespread biological function including diverse cellular processes like cell migration, differentiation, and apoptosis, by regulating RNA stability and translation(24–27). A peculiarity of *Drosophila* Unr is its dual sex specific function during dosage compensation. Contrary to its involvement in males, where it promotes the assembly of the MSL complex and thus is part of transcription regulation at least indirectly(28), it is involved in the inhibition of the same complex formation in female flies via translation repression of *msl2* mRNA, which is the rate limiting factor of the MSL complex(29, 30). Together with sex-lethal (Sxl), heterogeneous nuclear ribonucleoprotein 48 (Hrp48) and pAbp, Unr binds to the 3’ UTR and thereby inhibits the recruitment of the 43S pre-initiation complex(31–34). Structural details about assembly and action of this translation initiation repressor complex are scarce.

To conduct these widespread biological functions, Unr must interact with other RBPs, acting as a protein-RNA hub, that brings binding partners together and stabilizes their interaction(35). One example is its ternary interaction with Sxl and *msl2* mRNA during translation repression (29, 30, 36). Another well-established interactor is the poly(A)-binding protein (pAbp) (34, 37), which promotes translation upregulation and protection against mRNA decay through “closed-loop” formation of target mRNAs(38–42). However, in complex with Unr the fate of pAbp target mRNAs relies on the further composition of the RNP complex. On the one hand both proteins increase the mRNA stability of *c-fos* in complex with PAIP-1, hnRNP D and NSAP1 (43), whereas on the other hand it downregulates translation when accompanied with Imp1 on *pAbp* mRNA (44, 45).

Previous structural work on Unr focused on the first CSD and its RNA interaction or on single or multidomain constructs in unbound protein states (22, 36, 46). However, the role of the C-terminal CSDs has not been studied structurally, apart from showing that the last three CSDs bind RNA (22). Here we provide a high-resolution structure of a multidomain construct comprising these last three CSDs, showing for the first time the complexity of Unr-RNA interactions, which goes beyond the classical π-π-stacking of canonical CSDs. The relevance of the interaction was validated in several experiments and increases the knowledge about synergistic RNA binding which may occur in many additional multidomain RBPs. Moreover, we could identify interactions between a surface within the same C-terminal region and additional N-terminal regions of Unr with the *Drosophila* poly(A)-binding protein (pAbp). This gives atomistic insights into possible arrangements of Unr in a larger RNP translation regulatory complex, where it could rearrange the mRNA conformation and act as a hub for mRNP complex assembly.

## MATERIAL AND METHODS

### Plasmids

A pETM11 derived plasmid with a His_6_-affinity tag connected via a tobacco etch virus protease (TEV)-cleavage site to the protein constructs (derived from pBR322; G. Stier) was used for all protein expressions. Constructs of *Drosophila* Unr full length, CSD789 (A756-D990), CSD78 (A756-K922), ncCSD8 (P840-K922) and CSD9 (G911-D990) were used as described earlier(22). The different pAbp constructs, pAbp full length, pAbp RRM1 (A2-L84), pAbp RRM2 (G90-G176), pAbp RRM3 (G176-A263), pAbp RRM4 (L276-A362), pAbp linker (A362-N561) and pAbp PABC domain (K550-N634), were cloned from SL2 cDNA using the restriction free cloning approach. Point mutations for mutational analyses were inserted using site directed mutagenesis(47).

### Protein expression and purification

Protein expression and purification was done as described earlier(22). In brief, the proteins were expressed in *E. coli* BL21 (DE3) cells (*E. coli* B dcm ompT hsdS(r_B_^-^m_B_^-^) gal) using TB or isotope labeled M9 minimal medium, supplemented with ^15^NH^4^Cl and when needed ^13^C-D-glucose as sole nitrogen and carbon sources (purchased from Cambridge Isotope Laboratories). The cells were grown at 37°C before they were induced with 0.2 mM IPTG at an OD_600_ of 1.2 for TB and 0.8 for M9 minimal medium and incubated at 17°C for 16 h.

For protein purification the harvested cells were resuspended in 50 mM Hepes/NaOH pH 8.0, 500 mM NaCl, 30 mM imidazole, 1.4 mM b-mercaptoethanol and 1 M Urea and lysed using a French press. An affinity chromatography of the cleared lysate was done using 3 ml Ni-NTA gravity flow columns and the protein was eluted with 500 mM imidazole after an extensive wash with the lysis buffer. The proteins were cleaved using His_6_-TEV-protease and dialyzed against a low salt buffer (50 mM NaCl) without imidazole over night at 4°C. Cleavage for pAbp RRM2 was not successful, so that the purification was continued, and the samples were measured including the His_6_-tag. After a second nickel affinity purification, which gets rid of the protease and the cleaved tag, the protein was concentrated and injected on an S75 gel-filtration column (GE) for further purification and buffer exchange (20 mM NaP (pH 6.5), 50 mM NaCl, 1 mM DTT). Protein quality and purity was assessed by Coomassie staining and protein quantity by using a NanoDrop or BCA assay kit for the different protein constructs.

### NMR spectroscopy

All samples for NMR were measured in presence of 10% D_2_O and 0.01% NaN_3_ at 298 K on Bruker Avance III NMR spectrometers with magnetic field strengths corresponding to proton Larmor frequencies of 600 MHz, 700 MHz or 800 MHz equipped with triple resonance gradient cryogenic probe heads (600 and 800 MHz), a room temperature triple resonance probe head (700 MHz) or a room temperature quadrupole resonance probe head (600 MHz).

Experiments for backbone assignments were acquired on ^13^C and ^15^N labeled samples using conventional triple resonance experiments (backbone: HNCO, HNCA, CBCA(CO)NH and HNCACB(48)). For pAbp RRM1 0.03 mM (due to decreased solubility during the purification), pAbp RRM2 0.3 mM, pAbp RRM3 0.5 mM, pAbp RRM4 0.7 mM, pAbp linker 0.05 mM and pAbp PABC 1 mM samples were measured. Apodization weighted sampling was used for the acquisition of all spectra(49). These were processed using NMRPipe(50) and assigned with the program Cara(51). The backbone assignments of Unr CSD78, CSD789 and CSD9 were taken from Hollmann et al. (BMRB codes: 34492, 28086 and 34498)(22).

For NMR-based titrations ^15^N (or ^15^N^2^H for CSD789) labeled protein at a concentration of 0.1 mM (0.06mM for the competitive interaction study) was titrated with various ratios against purchased RNA oligonucleotides (A5, A7, A8, A9, A15, C8 and U8; IDT) or unlabeled protein. A ^1^H,^15^N-HSQC spectrum was recorded for each titration point. The RNA stocks were highly concentrated to keep dilution effects as small as possible (10 mM). The dilution was considered for peak intensity analysis. For protein titrations two samples were prepared, to avoid dilution effects. For titration analysis, Sparky(52) was used and the chemical shift perturbations (ppm) (CSP) at a ratio of 1:2 were calculated according to: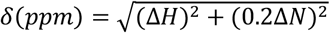(53). Shifts with a CSP greater than the average plus the standard deviation of all measured shifts were considered significant. The binding affinity reflected by the dissociation constant (K_d_) was obtained from a least square fit of the chemical shift changes for different residues during the titration, using: 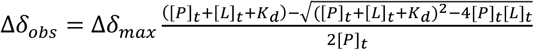(53), where x is the total ligand concentration and y is the corresponding CSP, P describes the protein concentration and A is the maximum CSP on saturation obtained as part of the fitting routine.

Standard pulse sequences were taken for the acquisition of *R*_*1*_, *R*_*2*_ and ^1^H-^15^N heteronuclear NOE experiments(54, 55) on a 0.1 mM ^15^N labeled deuterated sample. The relaxation delays were kept constant between all measurements (*R*_*1*_: 1600, 20, 50, 800, 100, 500, 150, 650, 1000, 400, 150 and 20 ms and *R*_*2*_: 25, 12.5, 50, 62.5, 100, 37.5, 75 and 25 ms). Peak integration and data fitting to derive spin relaxation parameters were done using PINT(56, 57). These parameters were taken to calculate the rotational correlation time (t_c_) according to Kay et al.(54) using: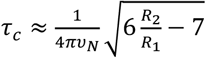

### Crystallography

A previously measured NMR sample of CSD789 bound to an A15-mer RNA was dialyzed against 10 liters of crystallization buffer (20 mM Hepes/NaOH pH 7.5, 50 mM NaCl and 1 mM DTT) to reduce the amount of phosphate, before being concentrated to 5 and 10 mg/ml. Several crystallization screens were set up at 7°C and 20°C and plate like crystals grew in 0.1 M Tris-Cl (pH 7.0), 0.2 M lithium sulfate and 2 M ammonium sulfate at 20°C. With a final size of 0.1×0.1×0.01 mm, these crystals were frozen in the mother liquor supplemented with 30 % glycerol and measured at the beamline P13 operated by EMBL Hamburg at the PETRA III storage ring (DESY Hamburg, Germany)(58). The crystal diffracted up to 1.2 Å and the data was processed in XDS(59). Molecular replacement was performed with CSD1 as search model (PDB: 4qqb)(36) using Phaser from Phenix suite(60, 61). Several rounds of model building in COOT(62) and refinement in the Phenix suite were done to further refine the structure.

For the crystal structure of Unr CSD789 bound to pAbp RRM3, crystals started to grow in an equimolar solution of both proteins and an A9-mer RNA oligonucleotide in 0.2 M sodium formate, 0.1 M bis-tris propane pH 6.5 and 20 % (w/v) PEG 3350 after two days. After three weeks the rectangular shaped crystals were frozen in the mother liquor supplemented with 40 % glycerol and measured at ID23-2 at the ESRF Grenoble. The crystal diffracted up to 2.9 Å. Data processing was done as described above, whereas the structure of CSD789 without RNA was used as search model for the molecular replacement. Data collection, processing, and refinement statistics for both structures are listed in Table S1.

### SAXS data acquisition and analysis

Small-angle X-ray scattering (SAXS) data were collected at the BioSAXS beamline BM29 at the ESRF, Grenoble,(63) using an X-ray wavelength of 0.992 Å and at the P12, operated by EMBL Hamburg at the Petra III storage ring (DESY Hamburg, Germany)(64) using an X-ray wavelength of 1.24 Å. 30 ml protein samples or buffer were purged through a quartz capillary for the measurements. Data acquisition details and statistics are listed according to community guidelines(65) in Table S2.

Prior analysis, frames were checked for radiation damage and then merged, and buffer subtracted. The data quality was checked by Guinier approximation. The whole analysis was done using the ATSAS 2.7.1 software package(66). CRYSOL(67) and EOM(68) calculations were done using the default settings to derive and fit theoretical scattering curves.

### Structure modeling

For the modeling of CSD789, high resolution structures of CSD78 (PDB: 6Y4H) and CSD9 (PDB: 6Y96)(22) were taken. The modeled structures were calculated using CNS (1.2)(69, 70) in an ARIA framework(71, 72). The structures were generated as described earlier(73). In brief, the single domains were connected to a single molecule, by implementing the missing linker residues. The linker region between CSD8 and CSD9 was randomized during the structure calculations. 5000 structures were calculated and fitted against a SAXS curve of CSD789.

To generate the model for the monomeric CSD789 15-mer poly(A)-RNA complex we first generated a pdb-file with the protein monomer and two 6-mer poly(A)-RNAs, where the first RNA is the molecule in the asymmetric unit and the coordinates for the second RNA were taken from the asymmetric unit adjacent to CSD9 in the unit cell. The residue numbering of the first RNA was kept fixed (1-6) while the second RNA was renumbered to follow the last residue of the first RNA (7-12) or with a gap of one to three additional adenosines (8-13, 9-14, 10-15). All missing residues of the protein including its hydrogen atoms and the RNA 15-mer were then generated in CNS-1.2(69) followed by an energy minimization step with fixed coordinates for the protein and the interacting RNA residues. Only the calculation where one adenosine was inserted in between the two RNA molecules and A6 was left flexible during the minimization gave a 15-mer RNA with proper geometry.

To generate the structural model in Figure 7b, we simply superimposed all available experimental high-resolution structures using PyMOL(74) (PDB accession codes: 4qqb, 6r5k, 6y6m, 6y6e, 6y4h, 6y96(22, 36, 75), and the structures determined in this study, PDB: 7zhh and 7zhr). The model of Unr-CSD1-9 was chosen from the SAXS-MD-based ensemble based on the proximity of ncCSD2 and ncCSD8 to allow interaction of both with pAbP-RRM12 and RRM3, respectively. A structural model of the Unr-ncCSD2-pAbP-RRM12 complex has been derived by modelling *Drosophila* pAbP-RRM12 in the RNA bound state using homology modelling (SWISS-MODEL(76)) based on the human pAbP-RRM12-RNA complex (PDB: 4F02(77)). To dock Unr-ncCSD2 with the pAbP-RRM12 model we used HADDOCK(78). Ambiguous interaction restraints to drive the docking were derived from chemical shift perturbations observed during NMR titrations between both proteins (Figure 5) and from their surface accessibility. These were for Unr-ncCSD2: T268, E269, T284, T285, R287, S289, C296 and F298; and for pAbP-RRM2: D102, K104, Y107, D108, S111, A112, G114 and N115. The default docking settings of the webserver were used. The resulting structure ensemble after water refinement consisting of 200 structures was clustered into 9 clusters, of which cluster 1 and 2 were by far the largest with 56 and 51 structures, respectively. Cluster 2 had the best statistics, with the best HADDOCK score and an average RMSD to the lowest energy structure of 0.63 ± 0.5 Å. The energy terms and buried surface area was also best for cluster 2. To incorporate this model into the structural model of the translation repression complex in Figure 7b, the pAbP-RRM domain containing structures (Unr-CSD789-pAbP-RRM3 and Unr-ncCSD2-pAbP-RRM12-RNA), were superimposed with the structure of yeast full-length PABP from PDB: 6R5K(79) to get an estimate about the distance of the RRM domains from each other in the RNA bound state.

### All-atom MD simulations of CSD789 and CSD789/RNA

The model of the CSD789/RNA complex was taken as starting conformation for all-atom molecular dynamics (MD) simulations. Simulations were carried out with the Gromacs software(80), version 2020.3. Interactions of the protein and the RNA were described with the Charmm36(81) force field, version March 2019, and the original TIP3P(82) water model was used. The protein and the protein-RNA complex were each placed in a dodecahedral box, where the distance between the protein to the box edges was at least 2.0 nm. The boxes were subsequently filled with water molecules and the systems were neutralized by adding sodium ions. In total, the systems contained at least 157761 atoms. The energy of the two systems were minimized within 400 ps with the steepest decent algorithm.

Subsequently, the systems were equilibrated for 100 ps with harmonic position restrains applied to the backbone atoms of the proteins (force constant 1000 kJ mol^-1^ nm^-2^). Finally, each simulation was simulated for 230 ns. The temperature was kept at 293 K using velocity rescaling (τ=0.1 ps)(83). The pressure was controlled at 1 bar with the Berendsen (τ =1 ps)(84) and with the Parrinello-Rahman barostat (τ=5 ps)(85) during equilibration and production simulations, respectively. The geometry of water molecules was constrained with the SETTLE algorithm(86), and LINCS(87) was used to constrain all other bond lengths. Hydrogen atoms were modeled as virtual sites, allowing a 4 fs time step. The Lennard-Jones potentials with a cut-off at 1nm were used to describe dispersive interactions and short-range repulsions. The pressure and the energy were corrected for missing dispersion interactions beyond the cutoff. Electrostatic interactions were computed with the smooth particle-mesh Ewald method(88, 89). Visual inspection of the simulations reveals that the RNA-protein contacts were stable throughout the simulation.

To ensure the agreement with NOE signals, distance restraints were applied in all simulations. In simulations of CSD789 with RNA, 20 distance restraints were applied. In simulations of only CSD789, 53 restraints were applied involving 258 atom pairs. Below a distance of 0.5 nm, no restraining potential was applied. Between 0.5 and 0.6 nm, a quadratic restraining potential with a force constant of 1000 kJ mol^-1^ nm^2^ was applied. Above 0.6 nm, a linear restraining potential was applied. The distance restraints are listed in Table S3. Distances that can contribute to a single NOE signal were treated simultaneously, implemented by defining with distances with the same restraint index in the Gromacs topology file.

### SAXS-driven MD simulation and SAXS calculations of CSD789 and CSD789/RNA

The SAXS-driven MD simulations and the subsequent SAXS calculations were performed with an in-house modification of Gromacs 2018.8, as also implemented by our webserver WAXSiS(90–92) for the SAXS calculations. The source code and documentation are available on GitLab at https://gitlab.com/cbjh/gromacs-swaxs and https://cbjh.gitlab.io/gromacs-swaxs-docs/, respectively. The simulation parameters were identical in MD and SAXS-driven MD simulations. Starting structures for the SAXS-driven simulations were taken from the last frame of the MD simulations. The SAXS restraints were turned on gradually over 15 ns and a force constant of 10 was used during the simulations. SAXS-restrained simulations were carried out for 50 ns. Simulation frames were saved every 2 ps for later analysis. Simulation frames from the time interval between 15 and 50 ns were used for the SAXS calculations. A spatial envelope was built around all solute frames of the protein and protein-RNA complex at a distance of 1.0 nm from all solute atoms. Because solvent atoms inside the envelope contributed to the SAXS calculations, the computed SAXS curves include effects from the hydration layer. The buffer subtraction was carried out using at least 351 simulation frames of a pure-water simulation box, which was simulated for 150 ns and which was large enough to enclose the envelope. The orientational average was carried out using 550 **q**-vectors for each absolute value of q, and the solvent electron density was corrected to the experimental value of 334 e/nm^3^, as described previously [12]. To compare the experimental with the calculated SAXS curve, we fitted the experimental curve via *I*_exp,fit_(*q*) = *f I*_exp_ + *c*, by minimizing chi-square with respect to the calculated curve. Here the factor *f* accounts for the overall scale, and the offset *c* takes the uncertainties from the buffer subtraction. No fitting parameters owing to the hydration layer or excluded solvent were used, implying that also the radius of gyration was not adjusted by the fitting parameters.

### MD simulations of CSD1–6 and CSD4–9

Ten different conformations for each CSD1–6 and CSD4–9 were taken from the rigid-body modelling and henceforth used as starting conformations for MD simulation. MD simulations were carried out with the Gromacs software, version 2019.6. Interactions of the protein were described with the Charmm36(81) force field, version March 2019, and the original TIP3P(82) water model was used. Each of the ten initial conformations was placed in a dodecahedral simulation box, where the distance between the protein to the box edges was at least 1.5 nm. The boxes with CSD1-6 were filled with 384,629 water molecules, and 1 chloride and 10 sodium ions were added to neutralize the systems. In total, these simulation systems contained 1,162,429 atoms. The boxes CSD4-9 were filled with 628,071 water molecules, and 1 chloride and 9 sodium ions were added to neutralize the systems. These simulation systems contained 1,893,654 atoms. The energy of each simulation system was minimized and equilibrated as described above for CSD789. Finally, each of the ten replicas was simulated for 230 ns without any restraints. The temperature was kept at 293 K using velocity rescaling (τ = 0.1 ps)(83). All other simulation parameters were set as described above for CSD789.

### SAXS calculations of CSD1–6 and CSD4–9

SAXS curves were computed from the free MD simulations using the same modified GROMACS version as described above for the CSD789. The distance between the protein and the envelope surface was at least 0.2 nm, such that nearly all water atoms of the hydration shell were included in nearly all frames. The buffer subtraction was carried out using 101 simulations frames taken from a 15 nm simulation of a pure-water simulation system, which was large enough to enclose the envelope. The orientational average was carried out using 14,200 *q*-vectors for CSD1–6 and 15,680 q-vectors for CSD4-9 for each absolute value of *q*. All other parameters and the fitting protocol were chosen as described above.

### Isothermal Titration Calorimetry

All titrations were done on a Malvern MicroCal PEAQ-ITC at 20°C while stirring at 750 rpm. The protein and RNA (IDT) samples were dialyzed against 20 mM NaP (pH 6.5), 50 mM NaCl and 0.5 mM TCEP. Concentrations of the molecules in each experiment are listed in Table S4. Each sample constellation was measured at least in duplicates and the MicroCal PEAQ-ITC analysis software was used to integrate, normalize, and fit the data.

### Fluorescence polarization assay

RNA oligonucleotides (AAA AAA AUG and A15-mer; IDT) were labeled at the 3’ end with fluorescein-5-thiosemicarbazide according to Qiu et al.(93). In brief, 5 μM RNA in 0.25 M sodium acetate (pH 5.6) were oxidized with 50 μM sodium periodate at 25°C in the dark for 90 minutes, before 100 μM of sodium sulfite were added. After 15 min at 25°C, 150 μM of fluorescein-5-thiosemicarbazide were added and the labeling reaction was performed for 3h at 37°C. The labeled RNA was precipitated for 3h at -80°C using one tenth of the reaction volume of 8 M LiCl and 2.5 times the reaction volume of 100 % ethanol. The concentration and labeling efficiency of the washed (70 % ethanol) and resuspended (H_2_O) pellet was measured using Nanodrop.

The fluorescence polarization assays were done in 20 mM NaP/NaOH pH 7.5, 50 mM NaCl and 2 mM DTT in a volume of 25 μl. 5 nM (AAA AAA AUG) or 25 nM (A15-mer) labeled RNA was incubated with different concentrations of protein for 30 min at 20°C. For each reaction a technical duplicate was measured in a black 384-well plate on a BioTek Synergy 4 plate reader using the corresponding filters and the automatic gain function. Each measurement was done in triplicates.

### Protein melting temperature

The nano differential scanning fluorimetry (nanoDSF) technology (nanotemper) was used to determine the protein melting temperature. Proteins were soaked into a standard capillary and heated up 1°C/min. The excitation varied from 10-30 % dependent on the protein concentration. The provided software was used to analyze the data and the melting temperature, at which 50 % of the protein was unfolded, was determined.

### Mass Photometry

The mass photometry analysis was performed using a Refeyn_MP_. The photometer was calibrated using filtered buffer (20mM sodium phosphate, 100mM NaCl). The proteins were diluted to a final concentration range between 5 and 60 nM. The measurement was immediately conducted after pipetting the protein on the sample carrier slide. To show the difference between immediate measurement and 10 minutes waiting time the sample was split into two, whereas one fraction was kept at room temperature for 10 minutes. Complex assembly was detected by change of back reflected scattering triggered by protein binding, which was monitored by the mass photometer. Assembled complexes resulted in an increased contrast through the scattering.

## RESULTS

### The C-terminal domain of Drosophila Unr tumbles independently, but with spatial restriction

Recently, we found that interdomain interactions between non-canonical CSDs and canonical CSDs of *Drosophila* Unr influence RNA target specificity (22). Robust domain-domain interactions could be found for CSD4-5, CSD5-6, and CSD7-8. Detailed information on RNA binding and consequences on domain arrangements are lacking. As we found that the three C-terminal domains bind to A-rich sequences (22), we aimed here to perform a thorough investigation of their RNA recognition mode. First, we investigated whether there is an inter-domain interaction for the two C-terminal domains (CSD8-9) as found for CSD7-8 and recorded ^15^N relaxation data of a Unr CSD789 construct to characterize the dynamics of the protein on a residue level (Fig. 1a). These data confirm the joint tumbling between CSD7 and ncCSD8(22), but also indicate that CSD9 tumbles independently of CSD78 due to the significant difference of the apparent rotational correlation times for CSD78 (t_c_ = 18.3±1.2 ns) and CSD9 (14.5±1.8 ns) (Fig. 1b). The linker between ncCSD8 and CSD9 is only 4 residues long and although the domains tumble independently, a SAXS driven structure modeling and MD simulation indicate spatial restrictions for CSD9 with respect to CSD78 (Fig. 1c and Fig. S1). Transient interactions between the two parts are confirmed by NMR data, which show chemical shift perturbations (CSPs) in a region within CSD9 (S970-S980) when compared to the longer construct of CSD789 (Fig 1d).

**Figure 1.**
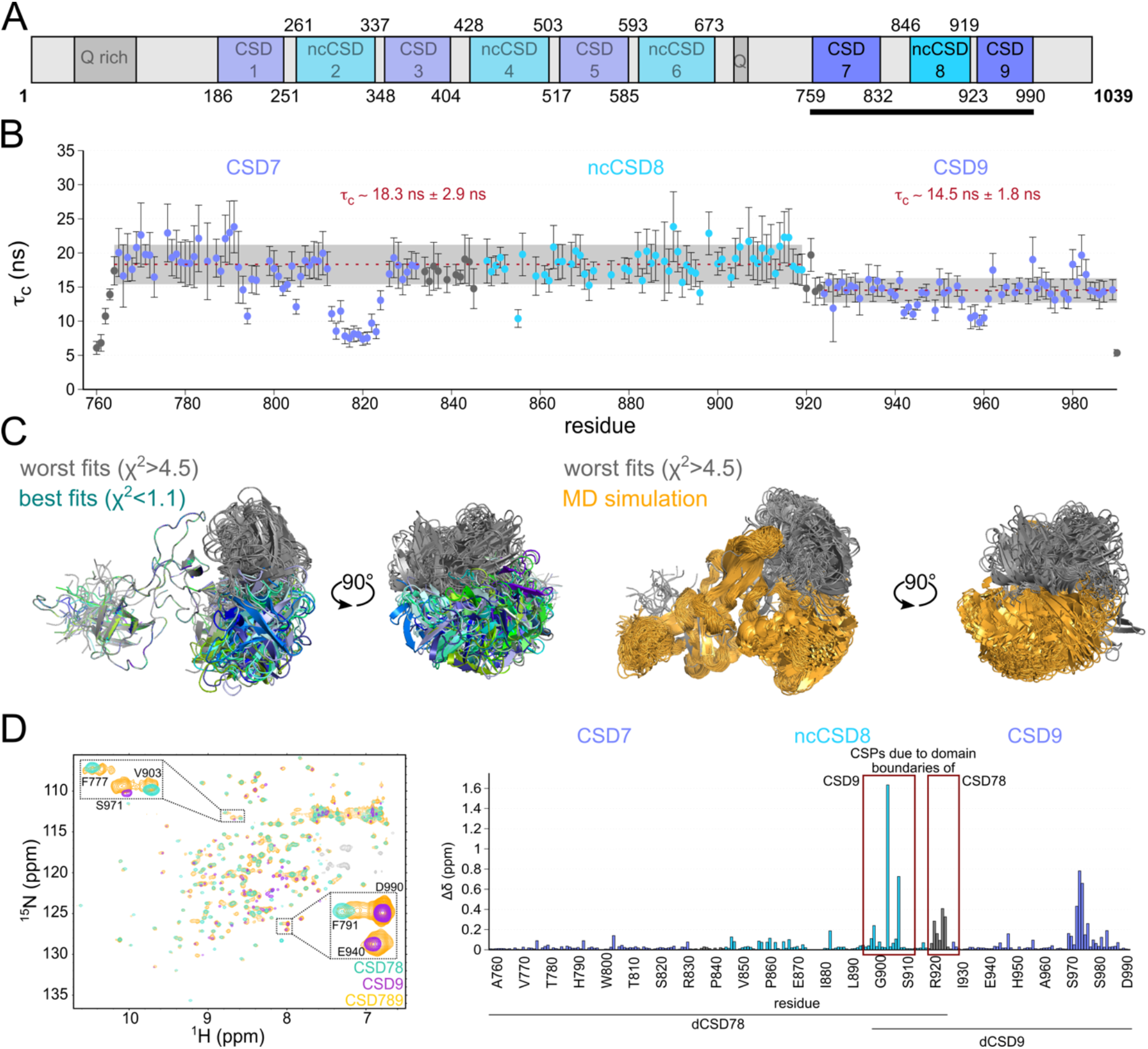
Restricted flexibility between ncCSD8 and CSD9 within CSD789 of *Drosophila* Unr. **a**: Schematic overview of CSD789 localization within the *Drosophila* Unr full length context. **b**: ^15^N relaxation data of CSD789 suggests that CSD9 tumbles independtly from CSD78 (CSD78: t_c_ = 18.3±2.9 ns vs. CSD9: t_c_ = 14.5±1.8 ns). The rotational correlation time (t_c_), which derives from the ^15^N transverse and longitudinal relaxation experiments, is plotted per residue. The error bars indicate the error propagation of the two relaxation experiments, which are initially derived from the quality of the exponential fit and the deviation between duplicates of two different relaxation delays. **c**: Structure modeling and a SAXS driven MD simualtion using SAXS data of Unr CSD789 indicate a restricted flexibility between ncCSD8 and CSD9. All structures are superimposed on CSD78 to see the flexibility of CSD9. Grey colored structures show the worst fitting structures, whereas green/blue show the best fitting ones. The MD simulated structures are shown in yellow. **d**: Overlaid ^1^H,^15^N-HSQC spectra of CSD78, CSD9 and CSD789 (left) showing CSPs in the termini of the single constructs (red boxes), but also within CSD9 (right). Canonical CSDs are colored blue throughout the whole figure and non-canonical CSDs cyan.

### Joint tumbling of the three C-terminal domains of Drosophila Unr upon RNA-binding

Next, we assessed RNA binding by CSD789, to examine whether this restricted flexibility between ncCSD8 and CSD9 has a similar importance on RNA binding and protein function as the fixed domain arrangement observed between the other domains. Previously strong binding of Unr CSD789 was shown for a poly(A)-15-mer RNA oligonucleotide(22), which is why poly(A) sequences of different length were used for this study.

Using NMR ^1^H,^15^N-HSQC titration experiments with poly(A) RNA sequences of different lengths (5-mer, 7-mer, 8-mer, 9-mer and 15-mer), a change between the binding affinities of the longer constructs compared to the shortest one (A5-mer) were observed (Fig. 2a-b and Fig. S2a-f). For some residues this change became clearly discernable by different binding modes. For the shorter A5-mer the CSPs induced by RNA binding are in the fast exchange regime^57^. In contrast, binding to RNA oligos with 7 adenines and longer result in an intermediate-to-slow exchange regime, indicative of a change in dissociation rates and therefore stronger binding (Fig. 2a and Fig. S2a-f). Not only the change of the exchange regime, but also the measured dissociation constants strengthen this observation. The affinity for the A5-mer is significantly lower than for the longer RNA oligonucleotides (Fig. 2b and Fig. S3a). To overcome the uncertainty of the affinity determination between CSD789 and the longer RNAs caused the change of the binding regime, isothermal titration calorimetry (ITC) was used for cross-validation. However, measurable thermal changes for the A5-mer could not be determined using ITC, potentially due to weak binding. Overall, the measured affinities using ITC are in line with the NMR observations (NMR: A5-mer: K_d_ = 127±16.6 μM vs. ITC: A7-mer: K_d_ = 17.7±5.2 μM, A8-mer: K_d_ = 4.4±2.7 μM, A9-mer: K_d_ = 4.3±1.3 μM and A15-mer: K_d_ = 4.8±0.8 μM). The non-gradual change in binding affinity and the saturation towards longer nucleotides strengthens the previously observed synergistic binding of the domains within CSD789^22^. A sufficient length of the RNA allows simultaneous binding of all domains.

**Figure 2.**
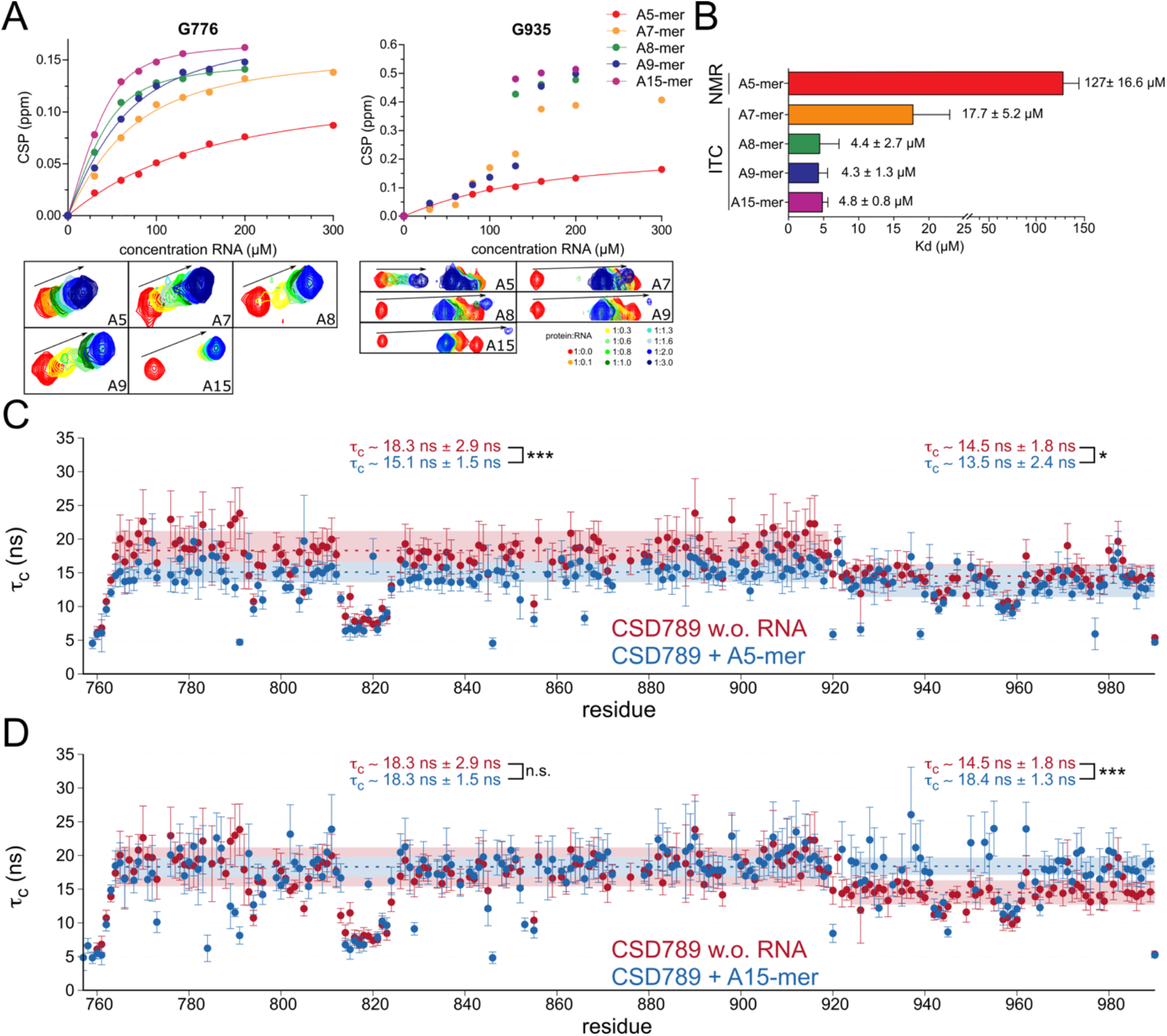
Joint tumbling of ncCSD8 and CSD9 within CSD789 upon RNA binding. **a**: NMR titrations and the corresponding fit of residues G776 and G935 of CSD789 titrations with poly(A)-mers of different length (A5-mer: red, A7-mer: orange, A8-mer: green, A9-mer blue, A15-mer purple). **b**: The averaged dissociation constants (K_d_), which were extracted from the NMR fit (A5-mer) or from at least three independent ITC measurements (A7-mer, A8-mer, A9-mer and A15-mer) are plotted for different RNA lengths. The error bars indicate the standard deviation. **c/d**: ^15^N relaxation data of CSD789 (Figure 1b) overlaid with data of the same protein bound to an A5-mer (**c**) (CSD78: t_c_ = 15.1±1.5 ns vs. CSD9: t_c_ = 13.5±2.4 ns) or an A15-mer (**d**) RNA (CSD78: t_c_ = 18.3±1.5 ns vs. CSD9: t_c_ = 18.4±1.3 ns) indicating, that CSD7-ncCSD8 and CSD9 only tumble together upon binding to the longer RNA. The rotational correlation time is plotted per residue. The error bars indicate the error propagation of the two relaxation experiments, which are initially derived from the quality of the exponential fit and the deviation between duplicates of two different relaxation delays. The statistical significance of differences in rotational correlation times was calculated using an unpaired t-test. N.s. meaning non significant, * meaning a p-value of P<0.05 and *** of P<0.001. Canonical CSDs are colored blue throughout the whole figure and non-canonical CSDs cyan.

Concomitantly, ^15^N relaxation data measured for the final titration point of the A15-mer RNA titration shows that the rotational correlation time of CSD9 increases to the overall tumbling time of CSD78 (CSD9: t_c_ = 14.5±1.8 ns unbound vs. 18.4±1.3 ns bound), indicating that the three domains tumble together in solution as one entity (Fig. 2d and Fig. S3c). In contrast, binding to the shorter A5-mer retains the independent tumbling of CSD9 towards CSD78, as observed for the unbound protein state (CSD9: t_c_ = 14.5±1.8 ns unbound vs. 13.5±2.4 ns bound) (Fig. 2c and Fig. S3b). The lower tumbling time of the domains CSD78 in the A5-mer bound state might reflect a reduced binding between these domains in an RNA bound form.

### CSD789 binding to poly(A) RNA sequence involves both typical and atypical CSD-RNA contacts

Due to the observed rigidification of Unr CSD789 upon binding to the longer RNA, we could successfully co-crystallize the protein with A15-mer RNA. The crystal structure could be solved using molecular replacement. From the 15 adenosines only six were visible in the structure, possibly due to flexibility of the remaining unbound nucleotides showing no electron density or due to degradation of the unbound nucleotides within the sample drop. Furthermore, one RNA chain is bound by two molecules from the same unit cell. CSD9 binds to RNA that is also bound by CSD7 and ncCSD8 of its symmetry mate, resulting in one unit cell having two protein and two RNA molecules (Fig. 3a). The signal-to-noise ratio of peaks in the NMR titration and rotational correlation times derived from ^15^N relaxation experiments show that despite high concentrations the complex has a 1:1 (protein:RNA) stoichiometry in solution. The rotation correlation time of around 18 ns fits what can be expected of the protein:RNA complex (around 15.7 ns at 295 K determined using ROTDIF(94)). The tumbling time for a 2:2 complex would be expected to be larger (Fig. 2d and Fig. S3c).

**Figure 3.**
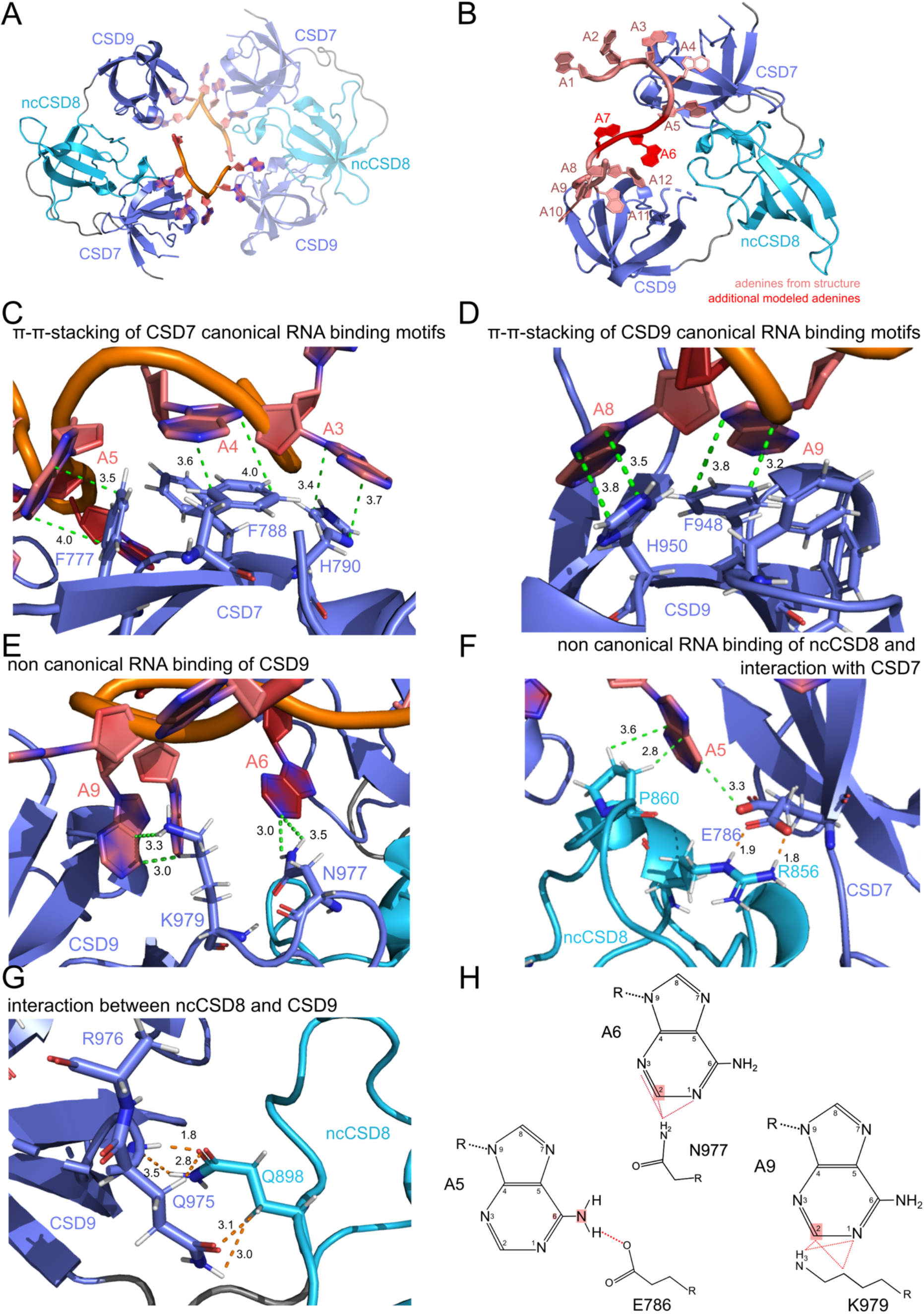
Crystal structure of CSD789 bound to a poly(A) RNA showing typical and atypical RNA contacts and domain-domain interactions. **a**: The crystal structure of CSD789 shows all three CSDs bound to five adenines of a poly(A) RNA (PDB: 7zhh). The symmetry mate in the same unit cell is highlighted by lower opacity. One RNA strand is bound to CSD7 and CSD9 of the symmetry mate. **b:** A data-driven model of a 1:1 CSD789-RNA complex in solution in which the two RNA strands from both symmetry mates (A1-A5 and A8-A12; pale red) are linked together by two additional nucleotides (A6 and A7; dark red) is shown. The protein-RNA contacts of the two additional nucleotides are validated by solution NMR and mutational analysis (see Fig. 4). **c/d:** The RNA binding residues within CSD7 (c; F777, F788 and H790) and CSD9 (d; F948 and H950) form the CSD typical p-p-stacking of the RNA bases and the aromatic rings. **e:** atypical RNA binding residues of CSD9 (N977 and K979) interact with the RNA base-specifically (A6 and A9). **f:** Residues of ncCSD8 (R856 and P860) interact with the RNA base-specifically (A5) and with E786 of CSD7. The electron density suggested two different conformations for E786, which are shown in the crystal structure. **g:** Interaction between two glutamines, one in ncCSD8 (Q868) and one in CSD9 (Q975), strengthen this domain-domain interaction. Dashed lines show the distance between atoms of an amino acid and a nucleotide (green) or between two amino acids (orange). The RNA is colored in red, canonical CSDs in blue and ncCSD8 in cyan. Due to potenital flexibility, no side chain could be build for R976. **h:** Adenine-specific interactions of CSD789 are illustrated. A hydrogen bond is formed between a free oxygen of E786 and NH6 of A5. Further interactions are formed between the hydrogens of C2 of A6 and A9 and the residues N977 and K979 respectively. The hydrogens have been added to the structure during modelling of the 1:1 complex. Canonical CSDs are colored blue throughout the whole figure and non-canonical CSDs cyan.

This suggests that the peculiar assembly seen in the crystal structure is a result of crystal packing. A structure that connects the termini of the single RNA strands by one additional nucleotide (A7) was modeled for a better impression of how the complex may look in solution (Fig. 3b and see methods). The validity of the model in solution is further confirmed by a SAXS driven molecular dynamics simulation (Fig. S2g) and by already described NMR titrations. The modeled nucleotide establishes contacts to residues in ncCSD8, of which corresponding NMR resonances shift significantly upon RNA titration (Fig. S2a-e). Interestingly the protein structure of human and *Drosophila* Unr CSD789 predicted by AlphaFold2(95), adopts the same domain orientations as our RNA-bound structure described here (Fig. S2h). Of note, the domain arrangement of CSD789 and its interdomain contacts are different from CSD456 (22).

As described previously for many other CSD structures(36, 96–102), the known RNA binding motifs (FGF and FFH) are involved in RNA binding of CSD7 and CSD9 (Fig. 3c, d). Surface exposed aromatic side chains of F777, F788 and H790 of CSD7 and F934, F948 and H950 of CSD9 form the p-p-stacking of the bases A3-A5 and A8-A9 of the RNA, resulting in a tightly packed interaction surface between the canonical CSDs and RNA. However, besides this typical RNA binding residues, previously unobserved atypical interactions are formed between N977 and A6 or K979 and A9 in CSD9. Strikingly these contacts form sequence-specific interactions towards adenine nucleotides (Fig 3e, h). Positioning of electronegative atoms to contact adenine-C2 would be sterically hindered by the exocyclic amino-N2 in guanine. Additionally, residues located in ncCSD8, that were already identified as RNA binding residues in NMR titration experiments (Fig. S2c-g)^22^ form direct contacts with nucleotide A5. The base points into the interaction surface of CSD7 and ncCSD8 and directly interacts with R856 and P860 (Fig. 3f). The arginine is additionally involved in an interaction with E786 of CSD7 and its electron density suggested two different confirmations within the structure (Fig. 3f, h). Due to hydrogen bonding of amino acids to the adenine-N1 or N6 adenine specific contacts are formed(103). Since E786 forms a hydrogen bond between its free oxygen to N6 of A5, the base gets sequence-specifically sandwiched between the two domains. To confirm the positional adenine specificity of CSD789 we performed NMR titrations of CSD789 with poly(U), poly(C), and poly(A) 8-mer RNAs (Fig. S4). A8-mer RNA addition induces large CSPs of CSD789 peaks already at substoichiometric concentrations, whereas poly(C) and poly(U) 8-mer RNAs induce weaker CSPs only at higher concentrations. Although we cannot presume that in certain positions other bases than an adenine would be preferred, we can confirm an overall adenine preference in solution.

The crystal structure further reveals domain-domain interactions, which are formed between glutamines located in ncCSD8 (Q898) and in CSD9 (Q975) that are pointing towards each other, and form hydrogen bonds as well as van der Waals contacts (Fig. 3g). The fact that the joint tumbling is strengthened in presence of RNA may be due to conformation changes upon RNA binding, which brings the two residues closer together. Further, this might also explain the restricted flexibility observed in the absence of RNA (Fig. 1c). Potentially there is a weak interaction between the two residues that is not strong enough to fix the two domains completely in absence of RNA but keeps them close in space through constant association and dissociation.

### Mutational analysis validates the solution model and confirms the importance of atypical RNA contacts and domain-domain interactions for RNA binding

To examine how much each of these interactions contribute to RNA binding and to validate the X-ray structure-based solution model, several mutants were tested for their binding affinity to an A15-mer RNA. Mutations were generated to disturb the atypical RNA binding of CSD9 (N977A and K979A), of ncCSD8 and its interface interactions to CSD7 (E786A, R856A and P860A) and the interactions between ncCSD8 and CSD9 (Q898A, Q975A and R976A). Electron density for the sidechain of R976 could not be detected. However, to exclude any kind of interaction between R976 with residues of ncCSD8 we included it in the mutational analysis (Fig. 4a). The mutated residues were in loop regions without secondary structure elements to avoid misfolding of the individual domains. High yield and solubility during the purification process, similar to wild type, peak dispersion in ^1^H,^15^N-HSQC spectra and the largely unchanged melting temperature demonstrates that the CSD fold for the mutants is not disrupted (Fig. 4b-c).

**Figure 4:**
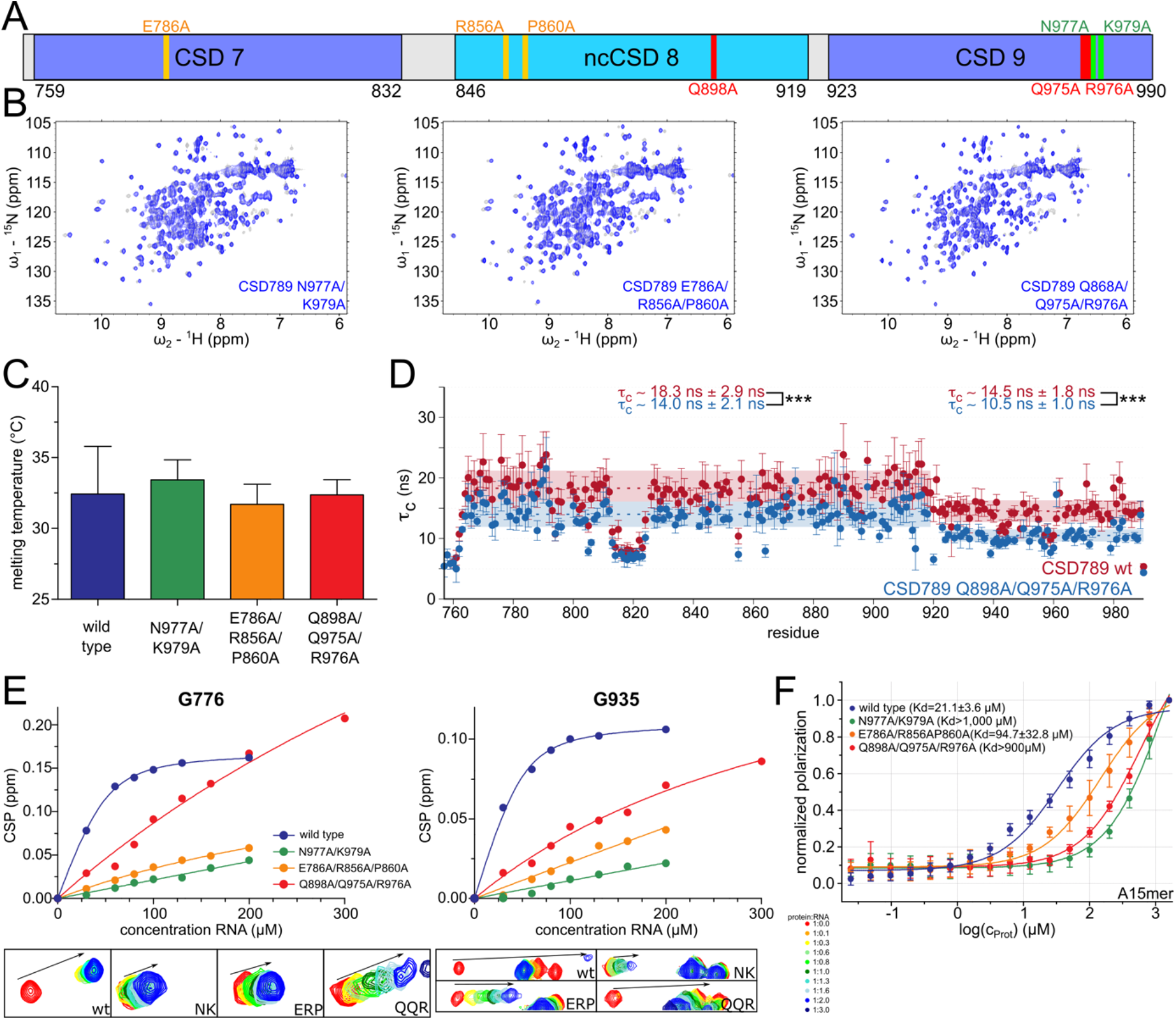
Mutational anaylsis of the solution structure of CSD789-A15-mer *in vitro*. **a**: The locations of mutations within CSD789 are highlighted schematically. **b:** ^1^H,^15^N-HSQC spectra of CSD789 wild type (grey) overlaid with the spectra of the differenet mutants (blue), indicating that mutations do not perturb the CSD789 fold. **c**: Melting temperatures for CSD789 wild type and the different mutants show no significant difference between the different constructs as determined by NanoDSF. Shown is the mean and standard deviation of three independent measurements. **d:** ^15^N relaxation data of CSD789 wild type protein (red; CSD78: t_c_ = 18.3±2.9 ns vs. CSD9: t_c_ = 14.5±1.8 ns) overlaid with data of a CSD789 Q898A/Q975A/R976A mutant (blue; CSD78: t_c_ = 14.0±2.1 ns vs. CSD9: t_c_ = 10.5±1.0 ns). The rotational correlation time is plotted per residue. The error bars indicate the error propagation of the two relaxation experiments, which are initially derived from the quality of the exponential fit and the deviation between duplicates of two different relaxation delays (see methods). The statistical significance of differences in rotational correlation times was calculated using an unpaired t-test. *** meaning a p-value of P<0.001. **e:** Chemical shift pertubations at different titration concentrations and the corresponding fit of two representative residues of the different CSD789 variants with a poly(A)-15-mer. **f:** Fluorescence polarization assays of CSD789 wild type and the different mutants to an A-15mer. Shown is the average binding affinity of at least three independent measurements with their standard deviation. The wildtype and mutant proteins are highlighted in different colors (wild type: blue, N977A/K979A: green, E786A/R856A/P860A: orange, Q898A/Q975A/R976A: red). Canonical CSDs are colored blue throughout the whole figure and non-canonical CSDs cyan.

**Figure 5.**
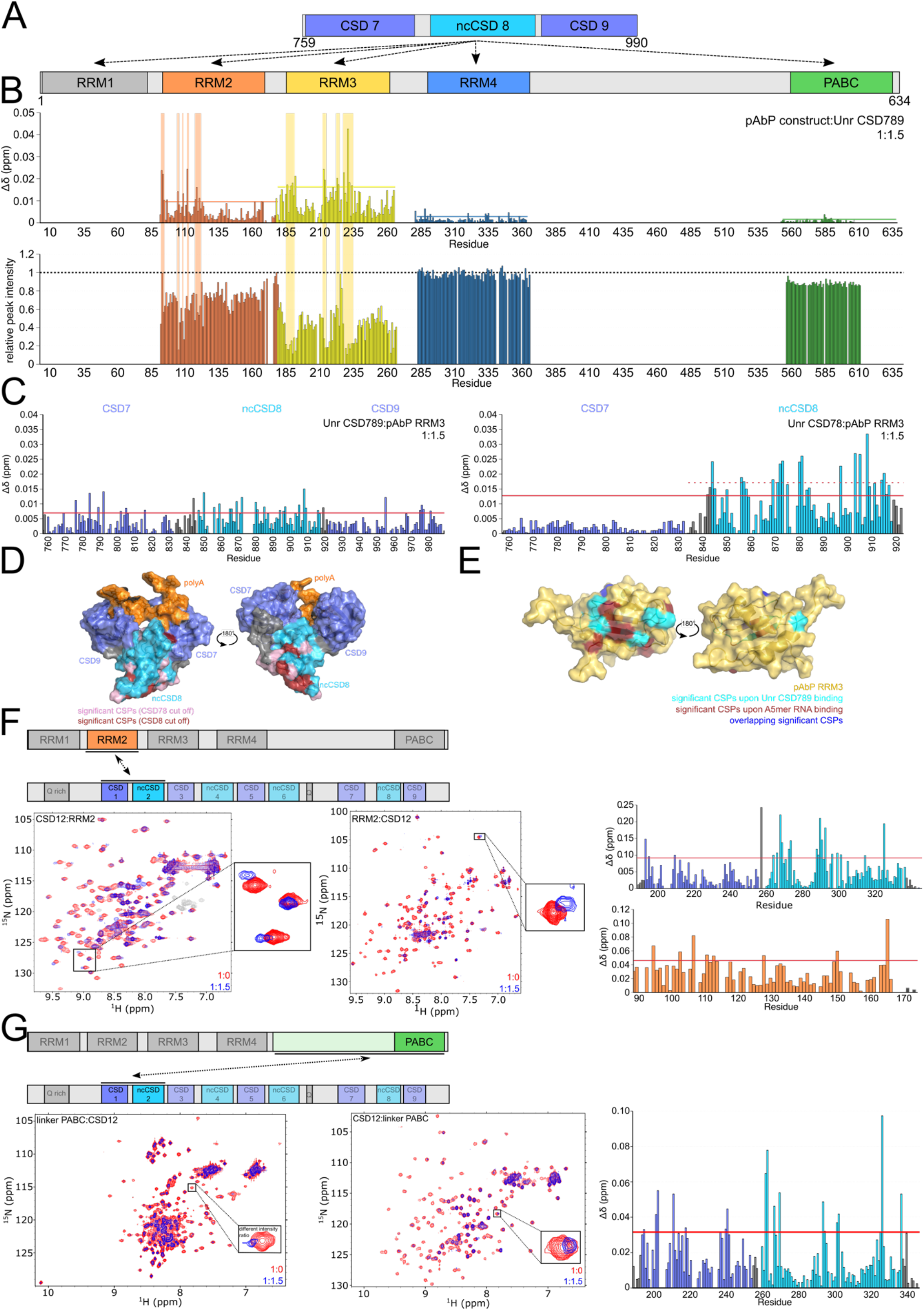
Unr interactions with the *Drosophila* poly(A)-binding protein (pAbp). **a**: Interaction of Unr CSD789 with the different domains within pAbp (RRM1-4 and PABC) was tested. **b:** CSP and intensity plots of differrent ^1^H,^15^N-HSQC NMR titration experiments with different ^15^N labeled pAbp constructs and unlabeled Unr CSD789. The red line in the CSP plots indicates the average plus standard deviation of all measured CSPs at a protein:protein ratio of 1:1.5, which was used to identify significant shifts(53). **c:** CSP plots of different ^1^H,^15^N-HSQC NMR titration experiments with ^15^N labeled Unr CSD789 and unlabeled pAbp RRM3 at a protein:protein ratio of 1:1.5. The red line in the CSP plots indicates the average plus standard deviation of all measured CSPs, which was used to identify significant shifts^57^. The dotted red line highlights the average plus standard deviation of the chemical shifts of CSD8 only. **d:** The significant shifts are lighlighted on the surface of the CSD789 structure bound to poly(A)-RNA. Residues of CSD78 with significant CSPs are highlighed in pink, whereas residues of CSD8 only with significant CSPs are colored in red. **e:** Residues with significant CSPs upon binding of CSD789 (cyan), an A5-mer RNA (red) and residues that overlap between the two titrations (blue) are highlighted on the surface of the pAbp RRM3 complex structure (yellow). **f/g**: ^1^H,^15^N-HSQC NMR spectra showing the interaction between Unr CSD12 and pAbp RRM2 (f) and the linker PABC region (g). The difference between the bound and unbound spectra are shown for each residue in the CSP plots. Canonical CSDs are colored blue throughout the whole figure and non-canonical CSDs cyan.

^15^N relaxation measurements of the Q868A/Q975A/R976A-mutant reveal lower rotational correlation times compared to the wild-type protein (CSD78: t_c_ = 14.0±2.1 ns mutant vs. 18.3±2.9 ns wild type; CSD9: t_c_ = 10.5±1.0 ns mutant vs. 14.5±1.8 ns wild type) (Fig. 4d and Fig. S5a). This indicates that the previously observed weak interaction between CSD78 and CSD9 is further weakened by these mutations. CSD78 and CSD9 are no longer temporarily interacting via the two glutamines, increasing the independent tumbling.

The effect of mutations on RNA binding was first assessed by NMR titration experiments. As saturation could not be reached, dissociation constants could not be reliably determined. This indicates that all mutants have a lower affinity to RNA compared to the wild type (Fig. 4e and Fig. S5b-e). The least impact on affinity had the Q868A/Q975A/R976A mutations, since chemical shift perturbations for e.g. G935 stayed in the intermediate exchange regime as observed for the wild type. For the E786A/R856A/P860A and N977A/K979A mutants the exchange regime changed from intermediate to fast exchange, indicative of weaker affinities. Concomitantly, fluorescence polarization (FP) data showed weaker binding for all mutants compared to the wildtype.

The binding curve of the wild-type protein results in a *K*_*d*_ of 21.1±3.6 μM to the A15-mer RNA (Fig. 4f). The measured *K*_*d*_ was lower compared to ITC (4.8±0.8 μM) or NMR (8.0±1.5 μM). The reason could be that the 3’ Cy5 label on RNA interferes with binding to CSD789. However, the *K*_*d*_ from our FP assays was similar to a previously reported affinity, which also used a Cy5 label in an EMSA (22). When comparing the binding affinity between wild type and mutants within the FP assay, all tested mutants show weaker binding to the two different RNAs. The mutant having a complete substitution of atypical RNA binding residues in ncCSD8 and its interface interactions to CSD7 (E786A/R856A/P860A) showed a more than fourfold weaker binding affinity with a *K*_*d*_ of around 94.7±32.8 μM compared to the wild type (Fig. 4f). Already single mutations showed a decreased binding affinity to A15-mer RNA (E786A: 50.1±6.4 μM; R856A: 72.4±11 μM; P860A: 67.2±7.9 μM) (Fig. S5f), meaning that each mutated amino acid contributes to RNA binding of CSD789.

The binding affinity of the other two interface mutants decreased even stronger. Here the *K*_*d*_ drops more than tenfold. Exact values cannot be calculated, since the binding does not even reach saturation with the maximum protein concentration range used (N977A/K979A: >1000 μM and Q868A/Q975A/R976A: >900 μM for both RNAs; Fig. 4f). As observed previously, also the single mutants bound the RNA significantly weaker than the wild-type protein (N977A: 71.2 ± 12.5 μM, K979A: 160.0 ± 57.6 μM, Q868A: 61.0 ± 12.1 μM, Q975A: 38.2 ± 8.7 μM and R976A: 161.0 ± 51.2 μM; Fig. S5f), indicating that all tested single mutations influence RNA binding.

Thus, all described structural peculiarities of CSD789, namely the atypical RNA binding within CSD9 and ncCSD8, the interface formation between CSD7 and ncCSD8 and ncCSD8 and CSD9 contribute to RNA binding of CSD789. The previously observed synergistic binding of CSD7 and CSD9 within CSD789 (Fig. 2a-d and Fig. S2a-f) could as such be explained by the solution structure presented here and is further validated by mutational analysis(22). Several atypical contacts contribute to RNA binding directly or indirectly via formation of additional domain-domain contacts. This validates our solution model of the CSD789-RNA structure and exemplifies the complexity of RNA binding by a multidomain RBP.

### Interaction of Unr with the poly(A)-binding protein pAbp

Our observations of sequence-specific binding of Unr CSD789 to adenines and a previously reported RNA independent interaction of *Drosophila* Unr with the *Drosophila* poly(A)-binding protein pAbp(22, 34, 37, 104), prompted us to understand the interaction between the two proteins in more detail. Firstly, we aimed to reproduce the interaction with recombinantly purified full-length proteins. Due to the low solubility of the full-length pAbp we have chosen to use mass photometry (Fig. S6a. Measurements for the individual proteins in solution show populations for the expected molecular weight (Unr: ∼90kDa vs. observed 95kDa; pAbp: ∼70kDa vs observed 60kDa). The pAbp sample shows additional populations (18% and 37%) of molecular weights of 199 and 440kDa likely due to aggregation, which would reflect the low solubility of full-length protein. However, the complex sample shows an additional peak at a molecular weight of 148kDa, corresponding to Unr-pAbp complex formation (calculated weight ∼ 160kDa).

To map the interaction sites of both proteins in more detail, we decided to perform extensive NMR chemical shift perturbation mapping by titrating 1.5 molar excess of Unr constructs (CSD12, CSD456 and CSD789) to single, isolated ^15^N-labelled pAbp domains (RRM1, RRM2, RRM3, RRM4 and the PABC domain) and record ^1^H,^15^N-HSQC experiments (Fig. 5a). Due to solubility problems, we could not perform the backbone assignments for RRM1.

A clear interaction could be identified between CSD789 and RRM2 and RRM3 (Fig. 5b and Fig. S6b). While the CSPs and intensity loss of RRM2 signals are less pronounced, which indicates that this interaction is potentially weak and unspecific, the CSPs, that form on two patches within the protein sequence of RRM3 are stronger and an overall intensity loss of around 50% is visible upon saturation with CSD789. The intensity of peaks that additionally show significant CSPs decrease more than 50%, indicative of an interaction between both proteins due to the increased size of the observed molecule and resulting slow molecular tumbling of the complex.

Similarly, a reverse titration of both proteins (^15^N labeled CSD789 and unlabeled RRM3) confirm this interaction (Figure 5c). However, a clear CSP pattern to enable mapping of the interaction site on CSD789 did not emerge. Therefore, we used shorter CSD constructs to narrow down the interaction site on CSD789. Since no significant shifts were traced for CSD7 or CSD9, this experiment identifies ncCSD8 as interaction partner of pAbp-RRM3 (Fig. 5c and Fig. S6c).

To get an impression of interaction sites between pAbp-RRM3 and CSD789, significant shifts of the CSD78 and of ncCSD8 titrations are mapped onto the available structures. The corresponding residues are mostly located on one site of the structure, on the opposite site of the RNA interaction surface of CSD789 and, identifying a clear surface for the interaction with pAbp RRM3 (Fig. 5d).

The reverse RNA titration shows that similar residues of RRM3, that are involved in RNA binding (Fig. S6d) are contributing to the interaction with CSD789. Significant CSPs of the interaction with CSD789 are located close to this RNA binding surface of the protein (Fig. 5e), suggesting that Unr competes with RNA for pAbp binding.

In addition to the ncCSD8-RRM3 interaction significant chemical shifts were observed between Unr CSD12 and pAbp RRM2 (Fig. 5f) and CSD12 and the linker PABC region of pAbp (Fig. 5g), indicative for additional protein interaction sites between both proteins.

We could then solve a crystal structure of Unr CSD789 bound to pAbp-RRM3 (Fig. 6a), which is consistent with the NMR data. As suspected pAbp-RRM3 binds to ncCSD8 of CSD789 with its RNA interaction surface. Thus, several amino acids of both interaction partners build up a large interaction platform (CSD789: E844, T845, H847, I871, E874, I880; RRM3: N184, I186, S212 and F227). The NMR data confirm a competitve binding between Unr CSD789 and the poly(A) RNA sequence. ^1^H-^15^N-HSQC data show especially for the RNA bound state of pAbp RRM3 large CSPs, whereas the binding to Unr CSD789 is mostly accompanied by intensity loss, mostly likely due to peak broadening because of the molecular weight increase. The spectrum in the presence of both ligands, shows a combination of both; signal loss on the one hand, but also stronger CSPs as associated with the RNA binding (Fig. 6b). These data suspect, that the RNA and Unr ncCSD8 are competing for the same interaction surface on pAbp-RRM3. Although we were not able to obtain a crystal structure of the ternary complex, a superposition of the two presented crystal structures shows that CSD789 would retain its capability to bind to RNA (Fig. 6c). To our knowledge, this is the first high-resolution structure of these two major translation regulatory proteins, Unr and pAbp in complex.

**Figure 6:**
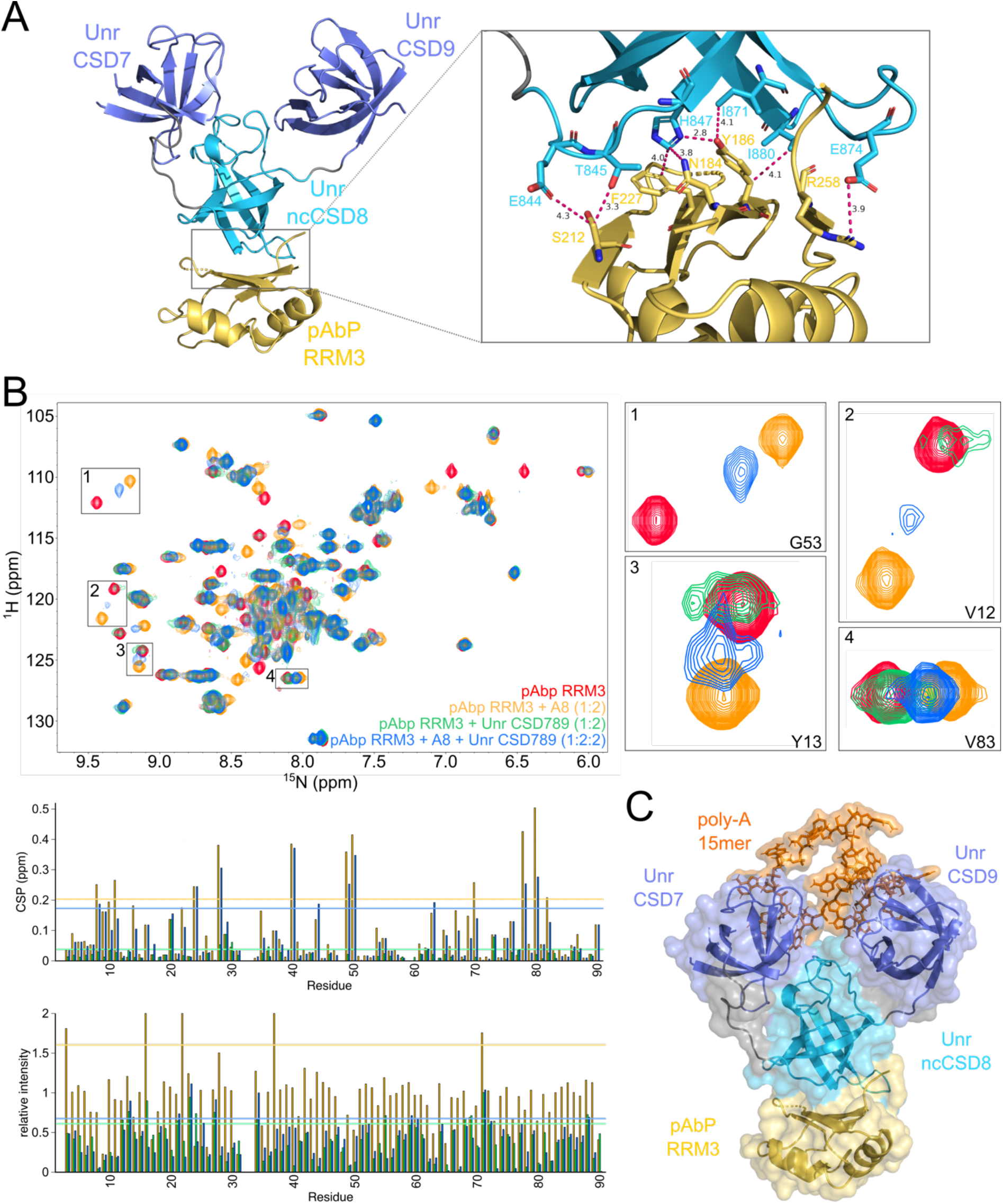
Structure determination of Unr CSD789 and pAbp RRM3. **a**: A crystal structure of Unr CSD789 (blue) and pAbp RRM3 (yellow) confirming the interaction surface observed by NMR titrations (PDB code: 7zhr). The detailed view on the interaction surface shows several amino acids of both proteins interacting with each other (right) (CSD789: E844, T845, H847, I871, E874, I880; RRM3: N184, Y186, S212 and F227). **b:** ^1^H,^15^N-HSQC NMR spectra showing the competitve binding between an A8-mer RNA and Unr CSD789 to pAbp RRM3. Shown is an overlay of spectra from apo pAbp RRM3 (red), bound to A8-mer (orange), titrated with Unr CSD789 (green) and titrated with both, A8-mer and Unr CSD789 (blue). Selected peaks were zoomed in to visualize the competition. The difference between the spectra are additionally illustrated for each residue in CSP and intensity ratio plots. **c:** A hybrid model generated by superposition of the crystal structure of CSD789 (blue) bound to a poly(A)-15mer (orange) and bound to pAbp RRM3 (yellow) visualizing that the interaction surfaces for both binding partners are on opposite sites. Canonical CSDs are colored blue and non-canonical CSDs in cyan.

**Figure 7:**
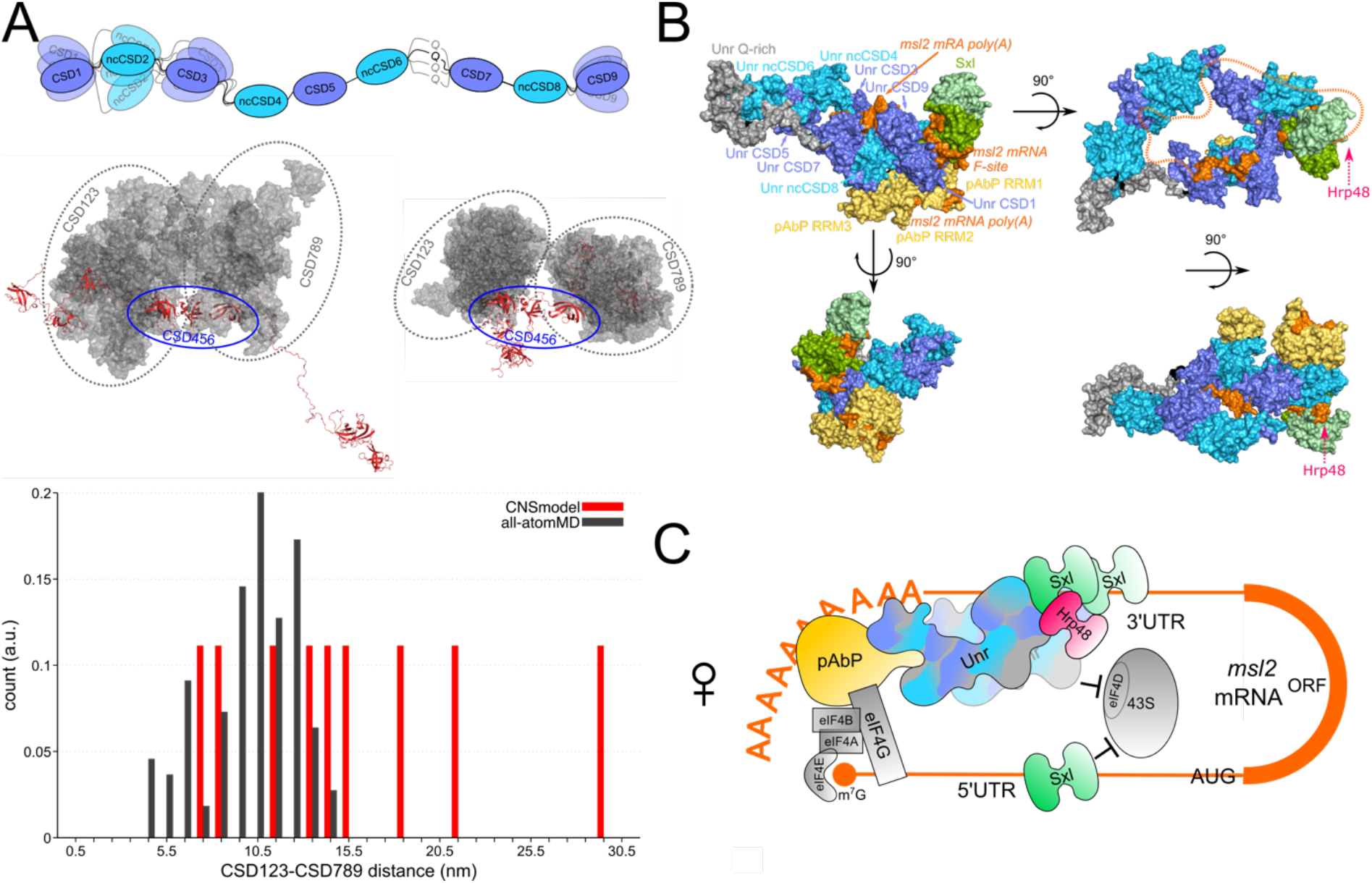
Structure modeling towards a deeper understanding of translation regulation. **a:** A schematic representation of the flexible regions within Unr, that were used for the almost full-length model generation (top). MD simulations showing that regardless of taking elongated or compact input structures (red), CSD123 and CSD789 come close in space (grey). All structures are superimposed on CSD456 (red and indicated with blue oval). Bar histogram, showing the distance distribution (nm) between CSD123 and CSD789 of the generated input structures (red) and the SAXS-supported MD simulations (grey). **b**: A structural model of the female dosage compensation complex in flies, derived by superposition of all available high resolution structures (PDB accession codes: 4qqb, 6r5k, 6y6m, 6y6e, 6y4h, 6y96(22, 36, 75), and the structures determined in this study, PDB: 7zhh and 7zhr). Additionally, a model of the Unr-CSD12-pAbP-RRM12 complex has been derived by data driven docking using HADDOCK(78) (see methods) based on chemical shift perturbations acquired in this study (Figure 5). A single model of Unr-CSD1-9 was chosen based on proximity of ncCSD2 and ncCSD8 which allow simultaneous interaction with pAbP-RRM2 and RRM3 respectively, and on absence of steric clashes. Unr is shown in blue, Sxl in green, pAbp in yellow and the RNA in green. The likely position of Hrp48 is indicated by a pink arrow. This model is solely for illustrational purposes to indicate that the identified interactions in all experimental high-resolution structures and NMR data can be consistent with a single structural model. **c:** A schematic representation of the complex shown in d. Additionally, the 5’ UTR binding site of Sxl and the repression of translation are highlighted. As opposed to the structural model in b), the full 3’ UTR translation repression complex likely features two Sxl and two Unr moieties due to the similarity of the E and F site of *msl-2* mRNA. Canonical CSDs are colored blue and non-canonical CSDs in cyan.

### MD simulations suggest a compact CSD1-9 conformation

RRM2 and 3 of pAbp are seperated only by a short linker (17 residues), suggesting that Unr CSD12, which interacts with pAbp RRM2, and Unr ncCSD8, which interacts with pAbp RRM3, are in close spatial proximity in the complex despite being spaced by 509 residues. To obtain the degree of extendedness or compactness of Unr, which would allow this interaction, SAXS data of longer Unr constructs were obtained. The SAXS data for CSD1-9 was affected by aggregation, so that datasets of CSD1-6 and CSD4-9 were used to validate MD simulations. Indeed, already free MD simulations of CSD1-6 and CSD4-9 show a good agreement with the SAXS data, suggesting that there is no need to further refine the simulations (Fig. S7).

Next, we modeled a CSD1–9 ensemble with correct bond geometries and without domain overlaps using CNS (1.2)(69, 70). The lack of flexible regions between several domains of Unr allowed us to generate model structures of almost full-length Unr, excluding the flexible N-terminal region, using the published high resolution structures of CSD12 (PDB: 6y6m), CSD456 (PDB: 6y6e), CSD78 (PDB: 6y4h) and CSD9 (PDB: 6y96)(105) and a homolgy model of CSD3(76, 106). The known rigid domain distances and orientations were kept fixed during the modelling. The remaining linker regions were randomized and thus allowed to adopt different conformations (Fig. 7a). The resulting structural ensemble covers a large conformational space involving center-of-mass (COM) distances between CSD123 and CSD789 up to 29 nm (Fig. 7a, red bars). To test if these conformations are compatible with the experiment-supported MD simulations, we superimposed the domains 4–6 from CSD1–6 and CSD4–9 MD-generated ensembles, thereby obtaining a plausible MD-based CSD1–9 ensemble (Fig. 7a). However, this MD-based ensemble was remarkably compact. Expanded conformations with COM > 15nm are not supported by the MD simualtions (Fig. 7a, black bars).

This suggests that some interdomain or domain-linker interactions restrict the overall flexibilty of the full-length protein, which is in accordance to the observed interactions of Unr with pAbp. This observation is of special interest considering that Unr is interacting directly with pAbp and Sxl in the female dosage compensation of flies. We could show in this study that Unr not only interacts with the F site region of *msl2* mRNA and Sxl(29, 30), but may also directly interact with the poly(A)-tail and pAbp.

## DISCUSSION

Although the number of RNA bound multidomain structures did increase within the last years (20, 21), atomistic insights into how multidomain RBDs exceeding two domains facilitate target recognition and specificity of RBPs is still scarce. Nevertheless, this knowledge is necessary to understand how full-length RBPs select for an RNA binding partner within the transcriptome. This study presents the first RNA bound multidomain structure of CSDs, indicating that their RNA binding mechanism can be far more complex than the previously shown π-π-stacking of the typical aromatic binding residues between a single CSD and RNA (Fig. 3)(102, 107, 108). Moreover, mutational interaction studies show that several atypical RNA binding residues contribute significantly to the RNA binding affinity of the C-terminal part of *Drosophila* Unr (Fig. 4). This structure not only increases our knowledge about the complex binding mechanism of multidomain RBD-RNA engagements, but it also suggests that the RNA binding of full-length Unr with its total five canonical and four ncCSDs is likely to be of even higher complexity and plasticity.

In an earlier study, we first speculated that CSPs upon RNA titration in ncCSD8 to be the result of unspecific interactions or allostery, as a single ncCSD8 construct did not harbor significant CSPs and a positively charged surface patch was located close to the interaction surface of CSD7(22). However, the crystal structure of CSD789 bound to the poly(A) RNA sequence showed that specific RNA-ncCSD8 contacts form and contribute to the overall binding affinity. Earlier identified binding motifs of Unr using SELEX, iCLIP or Shape analysis were rich of purine bases and especially adenosines(28, 108, 109), a phenomenon that cannot be sufficiently explained by the classical π-π-stacking of the canonical RNA binding motifs. The bacterial cold-shock proteins CspA and CspB, which only harbor a single CSD, were described as rather promiscuous RNA binders with low sequence specificity(110, 111) and CSD1 of Unr has low sequence specificity in isolation(36)..However, our structure shows a more complex interplay of multiple CSDs, where atypical interaction residues may be the main determinants for a specificity towards adenosines. Consequently, together with previous studies different mechanisms are shown to contribute to the target selectivity of Unr. As shown in this study, multiple CSDs increase the interaction surface of the RNA and the protein, allowing atypical binding residues to contribute to binding affinity and specificity of the protein. Additionally, a spatial restriction of the full-length protein, that gets introduced through interdomain contacts between canonical and non-canonical CSDs provides Unr with RNA tertiary structure specificity(22). A third mechanism to increase the target specificity of Unr is the interplay with additional RBPs, as shown in the case of the *msl2* mRNA, where interaction with Sxl is necessary to increase the binding affinity and base specificity of Unr CSD1 towards a certain cytosine within its target RNA sequence(36).

Considering this, the biological relevance of the poly(A) binding specificity of CSD789 gets strengthened with the identified interaction surface between ncCSD8 of Unr and RRM3 of pAbp. This characterized interaction validates previous pull-down interaction studies, that identified RRM3 of pAbp as the main driver for the interaction with Unr(37). The structure shows, that the interaction of both proteins blocks the RNA binding capability of RRM3 but keeps the binding site of CSD789 accessible for RNA. Therefore, it is possible that the C-terminal part of Unr binds close to the pAbp binding sites in the poly(A) tail or AU-rich elements. Since the main drivers of RNA interaction of pAbp are the first two RRMs(112, 113), an elimination of RRM3 from the RNA by CSD789 would not impact direct RNA binding of pAbp. Instead, RRM3 could be an important stabilizer of the Unr-CSD789-RNA interaction, by keeping the C-terminal part of Unr sandwiched between the RNA and itself. This keeps both proteins close together, which could increase the probability for additional interactions between them and potentiates RNA binding. We hypothesize, that this interaction is common during translation initiation of most mRNA targets and may promote recruitment of the 43S preinitiation complex. Indeed, it has been shown that a direct Unr-pAbp interaction stimulates tranlsation(104).

In the case of *msl2* translation repression, we propose that the presence of Sxl generates a conformational change within the Unr-mRNP complex by binding to CSD1, changing it from a general translation initiation promoter to a repressor complex. Here we present a more detailed model of how Unr could act as a molecular bridge, that ‘glues’ together the different components of the female dosage compensation complex in flies (Fig. 7b and c). This model has been generated by simple superposition of previously determined structures and structures presented in this study. A full-length Unr model from the SAXS-derived ensemble was chosen based on whether it allows the simultaneous binding of pAbp-RRM3 to ncCSD8 and pAbp-RRM2 binding to ncCSD2. This is only possible if both ncCSDs are in proximity by almost a circularized Unr. The 3’UTR of *msl2-*mRNA would wrap around Unr and Hrp48 is positioned just downstream of the F-site, which interacts with the 43S preinitiation complex and prevents it from binding to the 5’UTR.

Both presented high-resolution structures, showing the interaction of CSD789 with poly(A) RNA and with RRM3 of pAbp, contribute to a detailed structural insight of a larger translation regulation complex, which we hope to extend in the future. Unr and pAbp are identified to orchestrate translation initiation of different target mRNAs, whereby additional binding partners decide for the specific fate of the mRNA. Future studies will show, whether there are more interaction surfaces between both proteins, whether these are functionally relevant and whether the interaction between Unr and pAbp is invariable or dependent on the overall composition of this specific RNP.

## AVAILABILTY

The modified GROMACS software for SAXS-calculations and SAXS-driven MD simulations is available at https://gitlab.com/cbjh/gromacs-swaxs.

## ACCESSION NUMBERS

Structure coordinates have been deposited to the protein data bank (PDB) under the following accession codes: Unr CSD789 in complex with poly(A) RNA:7zhh, Unr CSD789 in complex with pAbp RRM3:7zhr. All NMR data have been deposited to the BMRB under the following accession codes (pAbp RRM2: 51392, pAbp RRM3: 51393, pAbp RRM4: 51394, pAbp PABC: 51395) and the SAXS data have been submitted to SASBDB (CSD789: SASDHH7, CSD1-6: SASDHM7, CSD4-9: SASDHL7).

## SUPPLEMENTARY DATA

Supplementary Data is available online.

## ACKNOWLEDGEMENT

We thank the ESRF Grenoble (beamlines BM29 and ID-30A) and DESY Hamburg PETRA-3 (P12 beamline) local contacts for support. We gratefully acknowledge Kathryn Perez and Karine Lapouge at the EMBL Protein Expression and Purification Facility for assisting with the ITC measurements.

## REFERENCES

1. Bieri, P., Greber, B.J. and Ban, N. (2018) High-resolution structures of mitochondrial ribosomes and their functional implications. Current Opinion in Structural Biology, 49, 44–53.

2. Voorhees, R.M. and Ramakrishnan, V. (2013) Structural Basis of the Translational Elongation Cycle. Annual Review of Biochemistry, 82, 203–236.

3. Sainsbury, S., Bernecky, C. and Cramer, P. (2015) Structural basis of transcription initiation by RNA polymerase II. Nature Reviews Molecular Cell Biology, 16, 129–143.

4. Cramer, P., Armache, K.-J., Baumli, S., Benkert, S., Brueckner, F., Buchen, C., Damsma, G.E., Dengl, S., Geiger, S.R., Jasiak, A.J., et al. (2008) Structure of Eukaryotic RNA Polymerases. Annual Review of Biophysics, 37, 337–352.

5. Lee, J. and Borukhov, S. (2016) Bacterial RNA polymerase-DNA interaction-The driving force of gene expression and the target for drug action. Frontiers in Molecular Biosciences, 3, 73.

6. Wilkinson, M.E., Charenton, C. and Nagai, K. (2020) RNA Splicing by the Spliceosome. Annual Review of Biochemistry, 89, 359–388.

7. Wan, R., Bai, R. and Shi, Y. (2019) Molecular choreography of pre-mRNA splicing by the spliceosome. Current Opinion in Structural Biology, 59, 124–133.

8. Wilkinson, M.E., Lin, P.-C., Plaschka, C. and Nagai, K. (2018) Cryo-EM Studies of Pre-mRNA Splicing: From Sample Preparation to Model Visualization. Annual Review of Biophysics, 47, 175–199.

9. Balcerak, A., Trebinska-Stryjewska, A., Konopinski, R., Wakula, M. and Grzybowska, E.A. (2019) RNA–protein interactions: disorder, moonlighting and junk contribute to eukaryotic complexity. Open Biology, 9.

10. Cech, T.R. and Steitz, J.A. (2014) The Noncoding RNA Revolution—Trashing Old Rules to Forge New Ones. Cell, 157, 77–94.

11. Gerstberger, S., Hafner, M. and Tuschl, T. (2014) A census of human RNA-binding proteins. Nature Reviews Genetics, 15, 829–845.

12. Hentze, M.W., Castello, A., Schwarzl, T. and Preiss, T. (2018) A brave new world of RNA-binding proteins. Nature Reviews Molecular Cell Biology, 19, 327–341.

13. Singh, G., Pratt, G., Yeo, G.W. and Moore, M.J. (2015) The Clothes Make the mRNA: Past and Present Trends in mRNP Fashion. Annual Review of Biochemistry, 84, 325–354.

14. Corley, M., Burns, M.C. and Yeo, G.W. (2020) How RNA-Binding Proteins Interact with RNA: Molecules and Mechanisms. Molecular Cell, 78, 9–29.

15. Auweter, S.D., Oberstrass, F.C. and Allain, F.H.-T. (2006) Sequence-specific binding of single-stranded RNA: is there a code for recognition? Nucleic acids research, 34, 4943–59.

16. Hennig, J. and Sattler, M. (2015) Deciphering the protein-RNA recognition code: Combining large-scale quantitative methods with structural biology. BioEssays, 37, 899–908.

17. Hennig, J., Gebauer, F. and Sattler, M. (2014) Breaking the protein-RNA recognition code. Cell cycle (Georgetown, Tex.), 13, 3619–20.

18. Cléry, A. and Allain, F.H.-T. (2011) RNA Binding Proteins (form structure to function of RNA binding proteins) - Madame Curie Bioscience Database.

19. Dimitrova-Paternoga, L., Jagtap, P.K.A., Chen, P.C. and Hennig, J. (2020) Integrative Structural Biology of Protein-RNA Complexes. Structure, 28, 6–28.

20. Schneider, T., Hung, L.-H., Aziz, M., Wilmen, A., Thaum, S., Wagner, J., Janowski, R., Müller, S., Schreiner, S., Friedhoff, P., et al. (2019) Combinatorial recognition of clustered RNA elements by the multidomain RNA-binding protein IMP3. Nature Communications, 10, 2266.

21. Gronland, G.R. and Ramos, A. (2017) The devil is in the domain: Understanding protein recognition of multiple RNA targets. Biochemical Society Transactions, 45, 1305–1311.

22. Hollmann, N.M., Jagtap, P.K.A., Masiewicz, P., Guitart, T., Simon, B., Provaznik, J., Stein, F., Haberkant, P., Sweetapple, L.J., Villacorta, L., et al. (2020) Pseudo-RNA-Binding Domains Mediate RNA Structure Specificity in Upstream of N-Ras. Cell Reports, 32, 107930.

23. Mihailovich, M., Wurth, L., Zambelli, F., Abaza, I., Militti, C., Mancuso, F.M., Roma, G., Pavesi, G. and Gebauer, F. (2012) Widespread generation of alternative UTRs contributes to sex-specific RNA binding by UNR. RNA, 18, 53–64.

24. Boussadia, O., Niepmann, M., Créancier, L., Prats, A.-C., Dautry, F. and Jacquemin-Sablon, H. (2003) Unr Is Required In Vivo for Efficient Initiation of Translation from the Internal Ribosome Entry Sites of both Rhinovirus and Poliovirus. Journal of Virology, 77, 3353–3359.

25. Dormoy-Raclet, V., Markovits, J., Jacquemin-Sablon, A. and Jacquemin-Sablon, H. (2005) Regulation of Unr expression by 5′- and 3′-untranslated regions of its mRNA through modulation of stability and IRES mediated translation. RNA Biology, 2, 112–120.

26. Elatmani, H., Dormoy-Raclet, V., Dubus, P., Dautry, F., Chazaud, C. and Jacquemin-Sablon, H. (2011) The RNA-Binding Protein Unr Prevents Mouse Embryonic Stem Cells Differentiation Toward the Primitive Endoderm Lineage. STEM CELLS, 29, 1504–1516.

27. Horos, R. and von Lindern, M. (2012) Molecular mechanisms of pathology and treatment in Diamond Blackfan Anaemia. British Journal of Haematology, 159, /a-n/a.

28. Militti, C., Maenner, S., Becker, P.B. and Gebauer, F. (2014) UNR facilitates the interaction of MLE with the lncRNA roX2 during Drosophila dosage compensation. Nature Communications, 5, 4762.

29. Duncan, K., Grskovic, M., Strein, C., Beckmann, K., Niggeweg, R., Abaza, I., Gebauer, F., Wilm, M. and Hentze, M.W. (2006) Sex-lethal imparts a sex-specific function to UNR by recruiting it to the msl-2 mRNA 3’ UTR: translational repression for dosage compensation. Genes & development, 20, 368–79.

30. Abaza, I., Coll, O., Patalano, S. and Gebauer, F. (2006) Drosophila UNR is required for translational repression of male-specific lethal 2 mRNA during regulation of X-chromosome dosage compensation. Genes & development, 20, 380–9.

31. Szostak, E., García-Beyaert, M., Guitart, T., Graindorge, A., Coll, O. and Gebauer, F. (2018) Hrp48 and eIF3d contribute to msl-2 mRNA translational repression. Nucleic acids research, 46, 4099–4113.

32. Beckmann, K., Grskovic, M., Gebauer, F. and Hentze, M.W. (2005) A dual inhibitory mechanism restricts msl-2 mRNA translation for dosage compensation in Drosophila. Cell, 122, 529–540.

33. Grskovic, M., Hentze, M.W. and Gebauer, F. (2003) A co-repressor assembly nucleated by Sex-lethal in the 3’UTR mediates translational control of Drosophila msl-2 mRNA. The EMBO Journal, 22, 5571–5581.

34. Duncan, K.E., Strein, C. and Hentze, M.W. (2009) The SXL-UNR Corepressor Complex Uses a PABP-Mediated Mechanism to Inhibit Ribosome Recruitment to msl-2 mRNA. Molecular Cell, 36, 571–582.

35. Guo, A.-X., Cui, J.-J., Wang, L.-Y. and Yin, J.-Y. (2020) The role of CSDE1 in translational reprogramming and human diseases. Cell Communication and Signaling, 18, 14.

36. Hennig, J., Militti, C., Popowicz, G.M., Wang, I., Sonntag, M., Geerlof, A., Gabel, F., Gebauer, F. and Sattler, M. (2014) Structural basis for the assembly of the Sxl–Unr translation regulatory complex. Nature, 515, 287–290.

37. Chang, T.C., Yamashita, A., Chen, C.Y.A., Yamashita, Y., Zhu, W., Durdan, S., Kahvejian, A., Sonenberg, N. and Shyu A. bin (2004) UNR, a new partner of poly(A)-binding protein, plays a key role in translationally coupled mRNA turnover mediated by the c-fos major coding-region determinant. Genes and Development, 18, 2010–2023.

38. Safaee, N., Kozlov, G., Noronha, A.M., Xie, J., Wilds, C.J. and Gehring, K. (2012) Interdomain Allostery Promotes Assembly of the Poly(A) mRNA Complex with PABP and eIF4G. Molecular Cell, 48, 375–386.

39. Wei, C.C., Balasta, M.L., Ren, J. and Goss, D.J. (1998) Wheat germ poly(A) binding protein enhances the binding affinity of eukaryotic initiation factor 4F and (iso)4F for cap analogues. Biochemistry, 37, 1910–1916.

40. Tarun, S.Z. and Sachs, A.B. (1996) Association of the yeast poly(A) tail binding protein with translation initiation factor eIF-4G. The EMBO Journal, 15, 7168–7177.

41. Wells, S.E., Hillner, P.E., Vale, R.D. and Sachs, A.B. (1998) Circularization of mRNA by eukaryotic translation initiation factors. Molecular Cell, 2, 135–140.

42. Goss, D.J. and Kleiman, F.E. (2013) Poly(A) binding proteins: Are they all created equal? Wiley Interdisciplinary Reviews: RNA, 4, 167–179.

43. Grosset, C., Chen, C.-Y.A., Xu, N., Sonenberg, N., Jacquemin-Sablon, H. and Shyu, A.-B. (2000) A Mechanism for Translationally Coupled mRNA Turnover: Interaction between the Poly(A) Tail and a c-fos RNA Coding Determinant via a Protein Complex. Cell, 103, 29–40.

44. Patel, G.P., Ma, S. and Bag, J. (2005) The autoregulatory translational control element of poly(A)-binding protein mRNA forms a heteromeric ribonucleoprotein complex.

45. Patel, G.P. and Bag, J. (2006) IMP1 interacts with poly(A)-binding protein (PABP) and the autoregulatory translational control element of PABP-mRNA through the KH III-IV domain. FEBS Journal, 273, 5678–5690.

46. Goroncy, A.K., Koshiba, S., Tochio, N., Tomizawa, T., Inoue, M., Watanabe, S., Harada, T., Tanaka, A., Ohara, O., Kigawa, T., et al. (2010) The NMR solution structures of the five constituent cold-shock domains (CSD) of the human UNR (upstream of N-ras) protein. Journal of Structural and Functional Genomics, 11, 181–188.

47. Braman, J., Papworth, C. and Greener, A. (1996) Site-Directed Mutagenesis Using Double-Stranded Plasmid DNA Templates. In In Vitro Mutagenesis Protocols. Humana Press, New Jersey, pp. 31–44.

48. Sattler, M., Schleucher, J. and Griesinger, C. (1999) Heteronuclear multidimensional NMR experiments for the structure determination of proteins in solution employing pulsed field gradients. Progress in Nuclear Magnetic Resonance Spectroscopy, 34, 93–158.

49. Simon, B. and Köstler, H. (2019) Improving the sensitivity of FT-NMR spectroscopy by apodization weighted sampling. Journal of Biomolecular NMR, 73, 155–165.

50. Delaglio, F., Grzesiek, S., Vuister, G.W., Zhu, G., Pfeifer, J. and Bax, A. (1995) NMRPipe: a multidimensional spectral processing system based on UNIX pipes. Journal of biomolecular NMR, 6, 277–93.

51. Keller, R. (2004) The Computer Aided Resonance Assignment tutorial. Goldau: CANTINA Verlag.

52. Lee, W., Tonelli, M. and Markley, J.L. (2015) NMRFAM-SPARKY: enhanced software for biomolecular NMR spectroscopy. Bioinformatics, 31, 1325–1327.

53. Williamson, M.P. (2013) Using chemical shift perturbation to characterise ligand binding. Progress in Nuclear Magnetic Resonance Spectroscopy, 73, 1–16.

54. Kay, L.E., Torchia, D.A. and Bax, A. (1989) Backbone Dynamics of Proteins As Studied by 15N Inverse Detected Heteronuclear NMR Spectroscopy: Application to Staphylococcal Nuclease. Biochemistry, 28, 8972–8979.

55. Zhu, G., Xia, Y., Nicholson, L.K. and Sze, K.H. (2000) Protein Dynamics Measurements by TROSY-Based NMR Experiments. Journal of Magnetic Resonance, 143, 423–426.

56. Niklasson, M., Otten, R., Ahlner, A., Andresen, C., Schlagnitweit, J., Petzold, K. and Lundström, P. (2017) Comprehensive analysis of NMR data using advanced line shape fitting. Journal of Biomolecular NMR, 69, 93–99.

57. Ahlner, A., Carlsson, M., Jonsson, B.H. and Lundström, P. (2013) PINT: A software for integration of peak volumes and extraction of relaxation rates. Journal of Biomolecular NMR, 56, 191–202.

58. Cianci, M., Bourenkov, G., Pompidor, G., Karpics, I., Kallio, J., Bento, I., Roessle, M., Cipriani, F., Fiedler, S. and Schneider, T.R. (2017) P13, the EMBL macromolecular crystallography beamline at the low-emittance PETRA III ring for high-and low-energy phasing with variable beam focusing. In Journal of Synchrotron Radiation. International Union of Crystallography, Vol. 24, pp. 323–332.

59. Kabsch, W., T., B.A., K., D., A., K.P., K., D., S., M., G., R.R.B., P., E., S., F., K., W., et al. (2010) XDS. Acta Crystallographica Section D Biological Crystallography, 66, 125–132.

60. Liebschner, D., Afonine, P. v., Baker, M.L., Bunkoczi, G., Chen, V.B., Croll, T.I., Hintze, B., Hung, L.W., Jain, S., McCoy, A.J., et al. (2019) Macromolecular structure determination using X-rays, neutrons and electrons: Recent developments in Phenix. Acta Crystallographica Section D: Structural Biology, 75, 861–877.

61. McCoy, A.J., Grosse-Kunstleve, R.W., Adams, P.D., Winn, M.D., Storoni, L.C. and Read, R.J. (2007) Phaser crystallographic software. urn:issn:0021-8898, 40, 658–674.

62. Emsley, P., Lohkamp, B., Scott, W.G. and Cowtan, K. (2010) Features and development of Coot. Acta Crystallographica Section D: Biological Crystallography, 66, 486–501.

63. Pernot, P., Round, A., Barrett, R., de Maria Antolinos, A., Gobbo, A., Gordon, E., Huet, J., Kieffer, J., Lentini, M., Mattenet, M., et al. (2013) Upgraded ESRF BM29 beamline for SAXS on macromolecules in solution. Journal of Synchrotron Radiation, 20, 660–664.

64. Blanchet, C.E., Spilotros, A., Schwemmer, F., Graewert, M.A., Kikhney, A., Jeffries, C.M., Franke, D., Mark, D., Zengerle, R., Cipriani, F., et al. (2015) Versatile sample environments and automation for biological solution X-ray scattering experiments at the P12 beamline (PETRA III, DESY). Journal of Applied Crystallography, 48, 431–443.

65. Trewhella, J., Duff, A.P., Durand, D., Gabel, F., Guss, J.M., Hendrickson, W.A., Hura, G.L., Jacques, D.A., Kirby, N.M., Kwan, A.H., et al. (2017) 2017 publication guidelines for structural modelling of small-angle scattering data from biomolecules in solution: An update. Acta Crystallographica Section D: Structural Biology, 73, 710–728.

66. Franke, D., Petoukhov, M. v, Konarev, P. v, Panjkovich, A., Tuukkanen, A., Mertens, H.D.T., Kikhney, A.G., Hajizadeh, N.R., Franklin, J.M., Jeffries, C.M., et al. (2017) ATSAS 2.8: a comprehensive data analysis suite for small-angle scattering from macromolecular solutions. Journal of applied crystallography, 50, 1212–1225.

67. Svergun, D., Barberato, C. and Koch, M.H.J. (1995) CRYSOL –a Program to Evaluate X-ray Solution Scattering of Biological Macromolecules from Atomic Coordinates. Journal of Applied Crystallography, 28, 768–773.

68. Tria, G., Mertens, H.D.T., Kachala, M. and Svergun, D.I. (2015) Advanced ensemble modelling of flexible macromolecules using X-ray solution scattering. IUCrJ, 2, 207–217.

69. Brunger, A.T. (2007) Version 1.2 of the Crystallography and NMR system. Nature Protocols, 2, 2728–2733.

70. Brünger, A.T., Adams, P.D., Clore, G.M., DeLano, W.L., Gros, P., Grosse-Kunstleve, R.W., Jiang, J.S., Kuszewski, J., Nilges, M., Pannu, N.S., et al. (1998) Crystallography & NMR system: A new software suite for macromolecular structure determination. Acta crystallographica. Section D, Biological crystallography, 54, 905–21.

71. Rieping, W., Habeck, M., Bardiaux, B., Bernard, A., Malliavin, T.E. and Nilges, M. (2007) ARIA2: Automated NOE assignment and data integration in NMR structure calculation. Bioinformatics, 23, 381–382.

72. Linge, J.P., Habeck, M., Rieping, W. and Nilges, M. (2003) ARIA: automated NOE assignment and NMR structure calculation. Bioinformatics, 19, 315–316.

73. Simon, B., Madl, T., Mackereth, C.D., Nilges, M. and Sattler, M. (2010) An Efficient Protocol for NMR-Spectroscopy-Based Structure Determination of Protein Complexes in Solution. Angewandte Chemie International Edition, 49, 1967–1970.

74. Wl, D. (2002) The PyMOL Molecular Graphics System. DeLano Scientific LLC, San Carlos, CA.

75. Schäfer, I.B., Yamashita, M., Schuller, J.M., Schüssler, S., Reichelt, P., Strauss, M. and Conti, E. (2019) Molecular Basis for poly(A) RNP Architecture and Recognition by the Pan2-Pan3 Deadenylase. Cell, 177, 1619-1631.e21.

76. Waterhouse, A., Bertoni, M., Bienert, S., Studer, G., Tauriello, G., Gumienny, R., Heer, F.T., de Beer, T.A.P., Rempfer, C., Bordoli, L., et al. (2018) SWISS-MODEL: homology modelling of protein structures and complexes. Nucleic Acids Research, 46, W296–W303.

77. Safaee, N., Kozlov, G., Noronha, A.M., Xie, J., Wilds, C.J. and Gehring, K. (2012) Interdomain Allostery Promotes Assembly of the Poly(A) mRNA Complex with PABP and eIF4G. Molecular Cell, 48, 375–386.

78. Dominguez, C., Boelens, R. and Bonvin, A.M.J.J. (2003) HADDOCK: A protein-protein docking approach based on biochemical or biophysical information. Journal of the American Chemical Society, 125, 1731–1737.

79. Schäfer, I.B., Yamashita, M., Schuller, J.M., Schüssler, S., Reichelt, P., Strauss, M. and Conti, E. (2019) Molecular Basis for poly(A) RNP Architecture and Recognition by the Pan2-Pan3 Deadenylase. Cell, 177, 1619-1631.e21.

80. Abraham, M.J., Murtola, T., Schulz, R., Páll, S., Smith, J.C., Hess, B. and Lindah, E. (2015) GROMACS: High performance molecular simulations through multi-level parallelism from laptops to supercomputers. SoftwareX, 1–2, 19–25.

81. Huang, J., Rauscher, S., Nawrocki, G., Ran, T., Feig, M., de Groot, B.L., Grubmüller, H. and MacKerell, A.D. (2016) CHARMM36m: an improved force field for folded and intrinsically disordered proteins. Nature Methods 2016 14:1, 14, 71–73.

82. Jorgensen, W.L., Chandrasekhar, J., Madura, J.D., Impey, R.W. and Klein, M.L. (1998) Comparison of simple potential functions for simulating liquid water. The Journal of Chemical Physics, 79, 926.

83. Bussi, G., Donadio, D. and Parrinello, M. (2007) Canonical sampling through velocity rescaling. The Journal of Chemical Physics, 126, 014101.

84. Berendsen, H.J.C., Postma, J.P.M., van Gunsteren, W.F., Dinola, A. and Haak, J.R. (1998) Molecular dynamics with coupling to an external bath. The Journal of Chemical Physics, 81, 3684.

85. Parrinello, M. and Rahman, A. (1998) Polymorphic transitions in single crystals: A new molecular dynamics method. Journal of Applied Physics, 52, 7182.

86. Miyamoto, S. and Kollman, P.A. (1992) Settle: An analytical version of the SHAKE and RATTLE algorithm for rigid water models. Journal of Computational Chemistry, 13, 952–962.

87. Hess, B. (2007) P-LINCS: A Parallel Linear Constraint Solver for Molecular Simulation. Journal of Chemical Theory and Computation, 4, 116–122.

88. Darden, T., York, D. and Pedersen, L. (1998) Particle mesh Ewald: An N·log(N) method for Ewald sums in large systems. The Journal of Chemical Physics, 98, 10089.

89. Essmann, U., Perera, L., Berkowitz, M.L., Darden, T., Lee, H. and Pedersen, L.G. (1998) A smooth particle mesh Ewald method. The Journal of Chemical Physics, 103, 8577.

90. Chen, P.C. and Hub, J.S. (2014) Validating solution ensembles from molecular dynamics simulation by wide-angle X-ray scattering data. Biophysical journal, 107, 435–447.

91. Chen, P.C. and Hub, J.S. (2015) Interpretation of Solution X-Ray Scattering by Explicit-Solvent Molecular Dynamics. Biophysical Journal, 108, 2573.

92. Knight, C.J. and Hub, J.S. (2015) WAXSiS: a web server for the calculation of SAXS/WAXS curves based on explicit-solvent molecular dynamics. Nucleic Acids Research, 43, W225.

93. Qiu, C., Liu, W.Y. and Xu, Y.Z. (2015) Fluorescence Labeling of Short RNA by Oxidation at the 3′-End. Methods in Molecular Biology, 1297, 113–120.

94. Berlin, K., Longhini, A., Dayie, T.K. and Fushman, D. (2013) Deriving Quantitative Dynamics Information for Proteins and RNAs using ROTDIF with a Graphical User Interface. Journal of biomolecular NMR, 57, 333.

95. Jumper, J., Evans, R., Pritzel, A., Green, T., Figurnov, M., Ronneberger, O., Tunyasuvunakool, K., Bates, R., Žídek, A., Potapenko, A., et al. (2021) Highly accurate protein structure prediction with AlphaFold. Nature 2021 596:7873, 596, 583–589.

96. Max, K.E.A., Zeeb, M., Bienert, R., Balbach, J. and Heinemann, U. (2007) Common mode of DNA binding to cold shock domains. FEBS Journal, 274, 1265–1279.

97. Sachs, R., Max, K.E.A., Heinemann, U. and Balbach, J. (2012) RNA single strands bind to a conserved surface of the major cold shock protein in crystals and solution. RNA (New York, N.Y.), 18, 65–76.

98. Yang, X.J., Zhu, H., Mu, S.R., Wei, W.J., Yuan, X., Wang, M., Liu, Y., Hui, J. and Ying Huang, X. (2019) Crystal structure of a Y-box binding protein 1 (YB-1)–RNA complex reveals key features and residues interacting with RNA. Journal of Biological Chemistry, 294, 10998–11010.

99. Yang, Y., Wang, L., Han, X., Yang, W.L., Zhang, M., Ma, H.L., Sun, B.F., Li, A., Xia, J., Chen, J., et al. (2019) RNA 5-Methylcytosine Facilitates the Maternal-to-Zygotic Transition by Preventing Maternal mRNA Decay. Molecular Cell, 75, 1188-1202.e11.

100. Zou, F., Tu, R., Duan, B., Yang, Z., Ping, Z., Song, X., Chen, S., Price, A., Li, H., Scott, A., et al. (2020) Drosophila YBX1 homolog YPS promotes ovarian germ line stem cell development by preferentially recognizing 5-methylcytosine RNAs. Proceedings of the National Academy of Sciences of the United States of America, 117, 3603–3609.

101. Welte, H., Zhou, T., Mihajlenko, X., Mayans, O. and Kovermann, M. (2020) What does fluorine do to a protein? Thermodynamic, and highly-resolved structural insights into fluorine-labelled variants of the cold shock protein. Scientific Reports, 10, 2640–2640.

102. Max, K.E.A., Zeeb, M., Bienert, R., Balbach, J. and Heinemann, U. (2006) T-rich DNA Single Strands Bind to a Preformed Site on the Bacterial Cold Shock Protein Bs-CspB. Journal of Molecular Biology, 360, 702–714.

103. Morozova, N., Allers, J., Myers, J. and Shamoo, Y. (2006) Protein-RNA interactions: exploring binding patterns with a three-dimensional superposition analysis of high resolution structures. Bioinformatics (Oxford, England), 22, 2746–52.

104. Ray, S. and Anderson, E.C. (2016) Stimulation of translation by human Unr requires cold shock domains 2 and 4, and correlates with poly(A) binding protein interaction. Scientific Reports, 6.

105. Ankush Jagtap, P.K., Müller, M., Masiewicz, P., von Bülow, S., Hollmann, N.M., Chen, P.-C., Simon, B., Thomae, A.W., Becker, P.B. and Hennig, J. (2019) Structure, dynamics and roX2-lncRNA binding of tandem double-stranded RNA binding domains dsRBD1,2 of Drosophila helicase Maleless. Nucleic Acids Research, 47, 4319–4333.

106. Guex, N., Peitsch, M.C. and Schwede, T. (2009) Automated comparative protein structure modeling with SWISS-MODEL and Swiss-PdbViewer: A historical perspective. ELECTROPHORESIS, 30, S162–S173.

107. Jacquemin-Sablon, H., Triqueneaux, G., Deschamps, S., le Maire, M., Doniger, J. and Dautry, F. (1994) Nucleic acid binding and intracellular localization of unr, a protein with five cold shock domains. Nucleic acids research, 22, 2643–50.

108. Triqueneaux, G., Velten, M., Franzon, P., Dautry, F. and Jacquemin-Sablon, H. (1999) RNA binding specificity of Unr, a protein with five cold shock domains. Nucleic acids research, 27, 1926–34.

109. Wurth, L., Papasaikas, P., Olmeda, D., Bley, N., Calvo, G.T., Guerrero, S., Cerezo-Wallis, D., Martinez-Useros, J., García-Fernández, M., Hüttelmaier, S., et al. (2016) UNR/CSDE1 Drives a Post-transcriptional Program to Promote Melanoma Invasion and Metastasis. Cancer Cell, 30, 694–707.

110. Graumann, P., Wendrich, T.M., Weber, M.H.W., Schröder, K. and Marahiel, M.A. (1997) A family of cold shock proteins in Bacillus subtilis is essential for cellular growth and for efficient protein synthesis at optimal and low temperatures. Molecular Microbiology, 25, 741–756.

111. Jiang, W., Hou, Y. and Inouye, M. (1997) CspA, the major cold-shock protein of Escherichia coli, is an RNA chaperone. The Journal of biological chemistry, 272, 196–202.

112. Sachs, A.B., Davis, R.W. and Kornberg, R.D. (1987) A single domain of yeast poly(A)-binding protein is necessary and sufficient for RNA binding and cell viability. Molecular and Cellular Biology, 7, 3268–3276.

113. Kühn, U. and Pieler, T. (1996) Xenopus poly(A) binding protein: Functional domains in RNA binding and protein-protein interaction. Journal of Molecular Biology, 256, 20–30.

