## Supplementary Figures and Tables for "Upstream of N-Ras C-terminal cold shock domains mediate poly(A) specificity in a novel RNA recognition mode and bind poly(A) binding protein during translation regulation"

**This PDF file includes:**

Figures S1-S7
Tables S1-S4

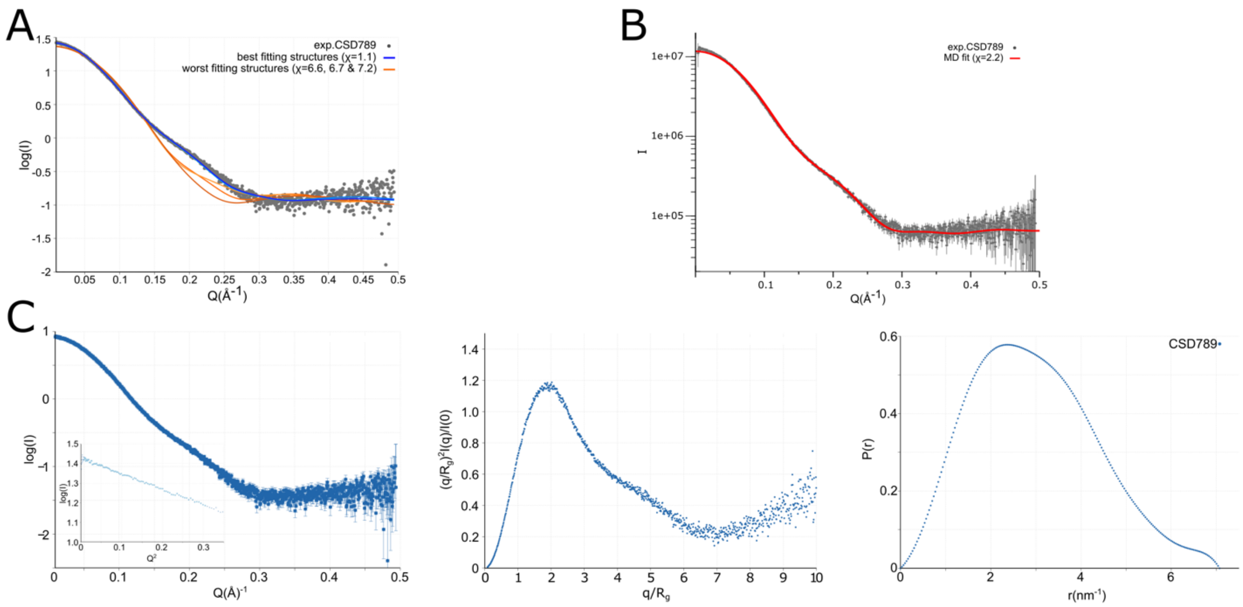


**Figure S1 a**: Crysol fit of six different structures of CSD789 of 1000 modeled structures to the experimentally measured SAXS curve of CSD789. The best fitting curves are highlighted in blue, whereas the worst fits are colored in orange. **b:** The fit of a SAXS driven MD simualtion plotted in the scatterig curve of the protein in solution. **c**: left: *I(q) versus q* as log-linear plots with the inset showing the Guinier fits for $qR_g$ < 1.3 indicating good data quality and no aggregation for the SAXS curve of CSD789. Middle: Dimensionless Kratky plots indicate that CSD789 contains low flexibility and is mostly structured. Right: Normalized *P(r) versus r* profile shows the radius of gyration for CSD789.

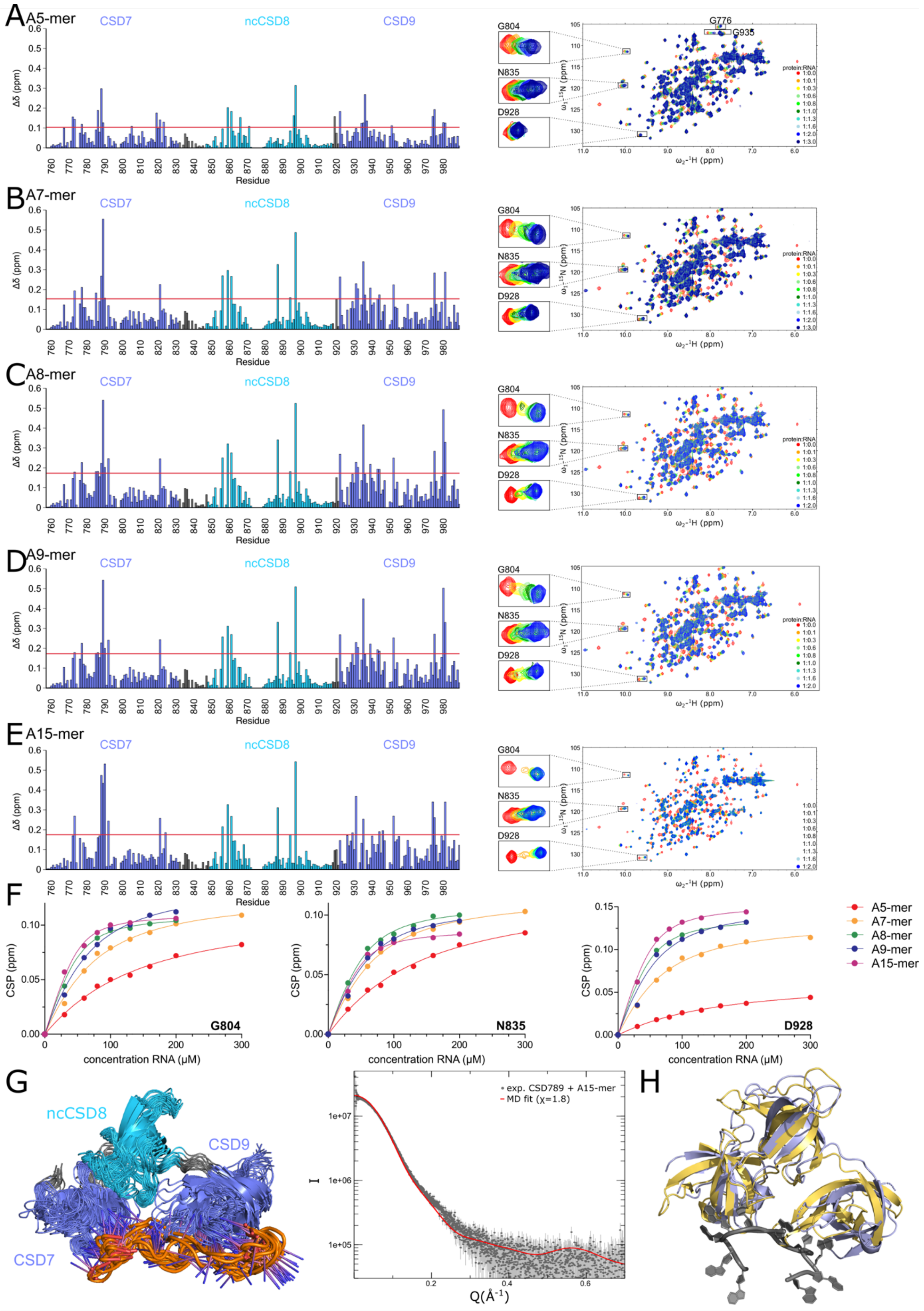
A
A5-mer
B
A7-mer
C
A8-mer
D
A9-mer
E
A15-mer
CSD7
ncCSD8
CSD9
Δδ (ppm)
Residue
G804
N835
D928
G776
G935
protein:RNA
ω₁-¹⁵N (ppm)
ω₂-¹H (ppm)
F
CSP (ppm)
concentration RNA (μM)
A5-mer
A7-mer
A8-mer
A9-mer
A15-mer
G
H
exp. CSD789 + A15-mer
MD fit (χ=1.8)
I
Q(Å⁻¹)

**Figure S2:** $^{1}H,^{15}N$-HSQC NMR titration experiments with CSD789 and the calculated chemical shift pertubations plots for an A5-mer (**a**), A7-mer (**b**), A8-mer (**c**), A9-mer (**d**) and A15-mer (**e**) at a protein:RNA ratio of 1:2. The red line in the CSP plots indicates the average plus the standard deviation of all measured shifts, which was used to identify significant shifts[58]. **f:** Corresponding fits of three different residues of CSD789 titrations with the different poly(A)-mers (A5-mer: red, A7-mer: orange, A8-mer: green, A9-mer blue, A15-mer purple). **g**: SAXS driven molecular dynamics simulation validating the strucutral solution model of Unr CSD789 bound to a 15-mer poly(A) RNA sequence. **h:** Comparison of the crystal structure of CSD789 bound to RNA (orange) and the predicted fold of the human CSD789 from AlphaFold (grey). Canonical CSDs are colored blue throughout the whole figure and non-canonical CSDs cyan.

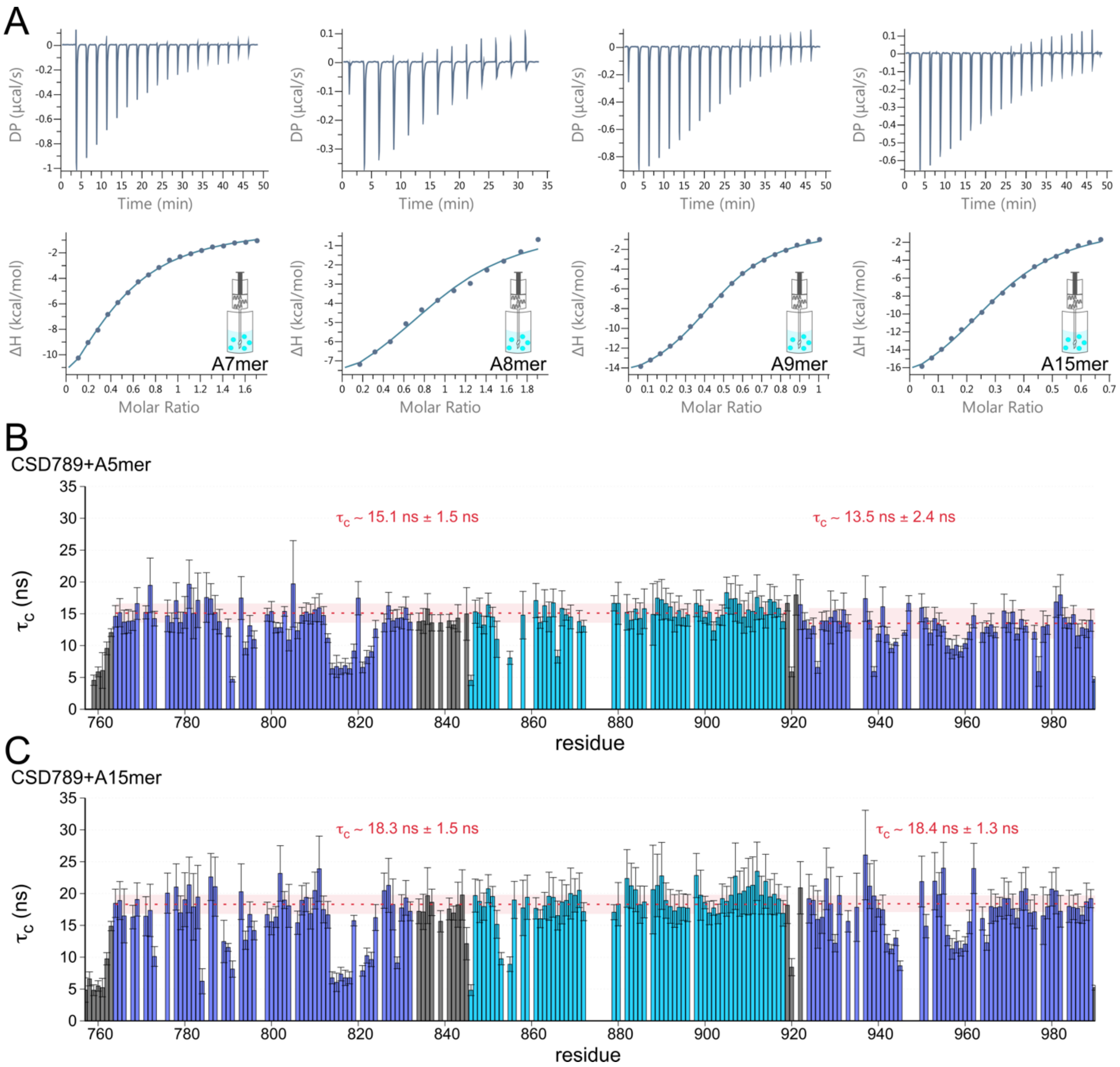


**Figure S3 a**: ITC calorimetric data for CSD789 binding to poly(A) RNA sequences of different length. The RNA was titrated from the syringe into the cell which contains the protein. The data are exemplary for one replicate of at least three independent measurements. **b/c:** $^{15}N$ relaxation data of CSD789 bound to A5-mer (**a**) or A15-mer (**b**) RNA. The rotational correlation time ($\tau_c$) derived from $^{15}N$ longitudinal and transverse relaxation experiments is plotted per residue. The error bars indicate the error propagation from errors of the two relaxation experiments, which are derived from the quality of the exponential fit and the deviation between duplicates of two different relaxation delays. Canonical CSDs are colored blue throughout the whole figure and non-canonical CSDs cyan.

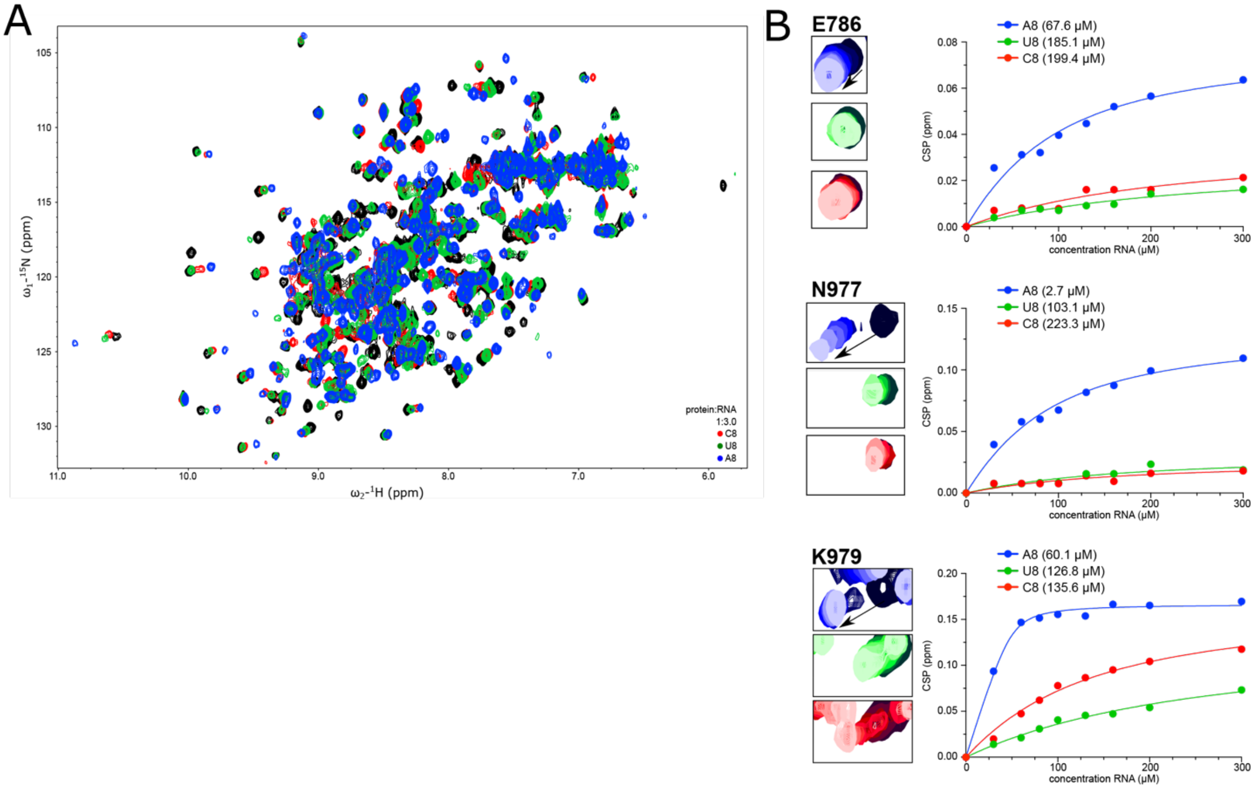


**Figure S4 a:** Full view of $^{1}$H,$^{15}$N-HSQC NMR spectra comparing the chemical shift perturbation between free CSD789 (black) and bound to poly(A) (blue), poly(U) (green) and poly(C) (red). **b:** zooms into spectral regions of peak corresponding to residues involved in adenine specfic RNA binding (E786, N977 and K979). At the right, binding curves are fitted to determine dissociation constants.

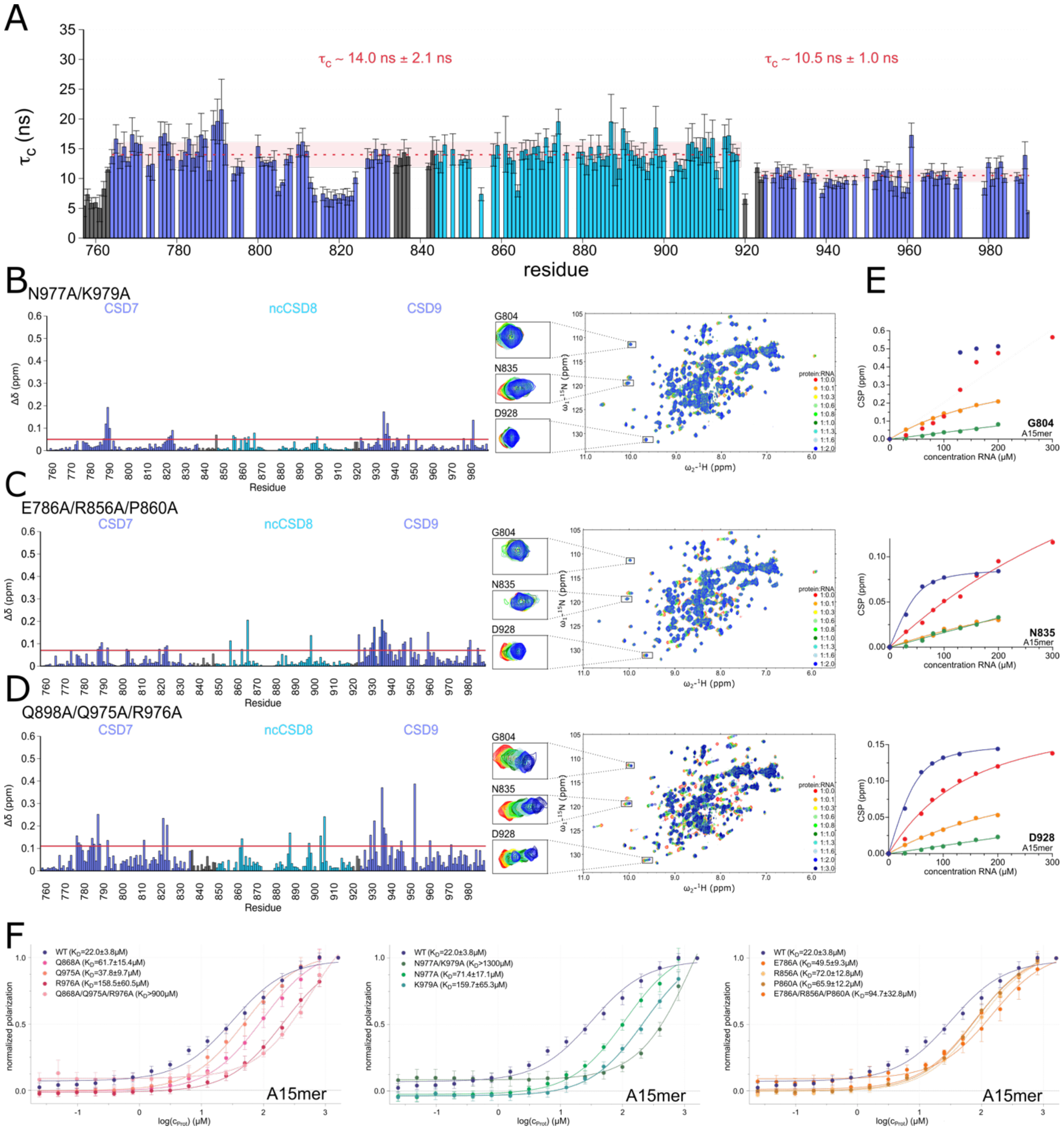


**Figure S5 a**: $^{15}$N relaxation data of CSD789 Q898A/Q975A/R976A. The rotational correlation time ($\tau_c$) derived from $^{15}$N longitudinal and transverse relaxation experiments is plotted per residue. The error bars indicate the error propagation from errors of the two relaxation experiments, which are derived from the quality of the exponential fit and the deviation between duplicates of two different relaxation delays. **b/c/d:** $^{1}$H,$^{15}$N-HSQC NMR titration experiments of the different CSD789 mutant constructs and the calculated chemical shift pertubation plots for an A15-mer binding at a protein:RNA ration of 1:2. The red line in the CSP plots indicates the average plus the standard deviation of all measured shifts, which was used to identify significant shifts[58]. **e:** Chemical shift pertubations at different titration concentrations and the corresponding fit of three different residues of the different CSD789 constructs with a poly(A)-15-mer. **f:** Fluorescence polarization assays of CSD789 wild type and the different mutants to an A15-mer RNA sequence, showing the different binding affinities between the constructs. Shown is the average of at least three independent measurements and the standard deviation. Canonical CSDs are colored blue throughout the whole figure and non-canonical CSDs cyan.

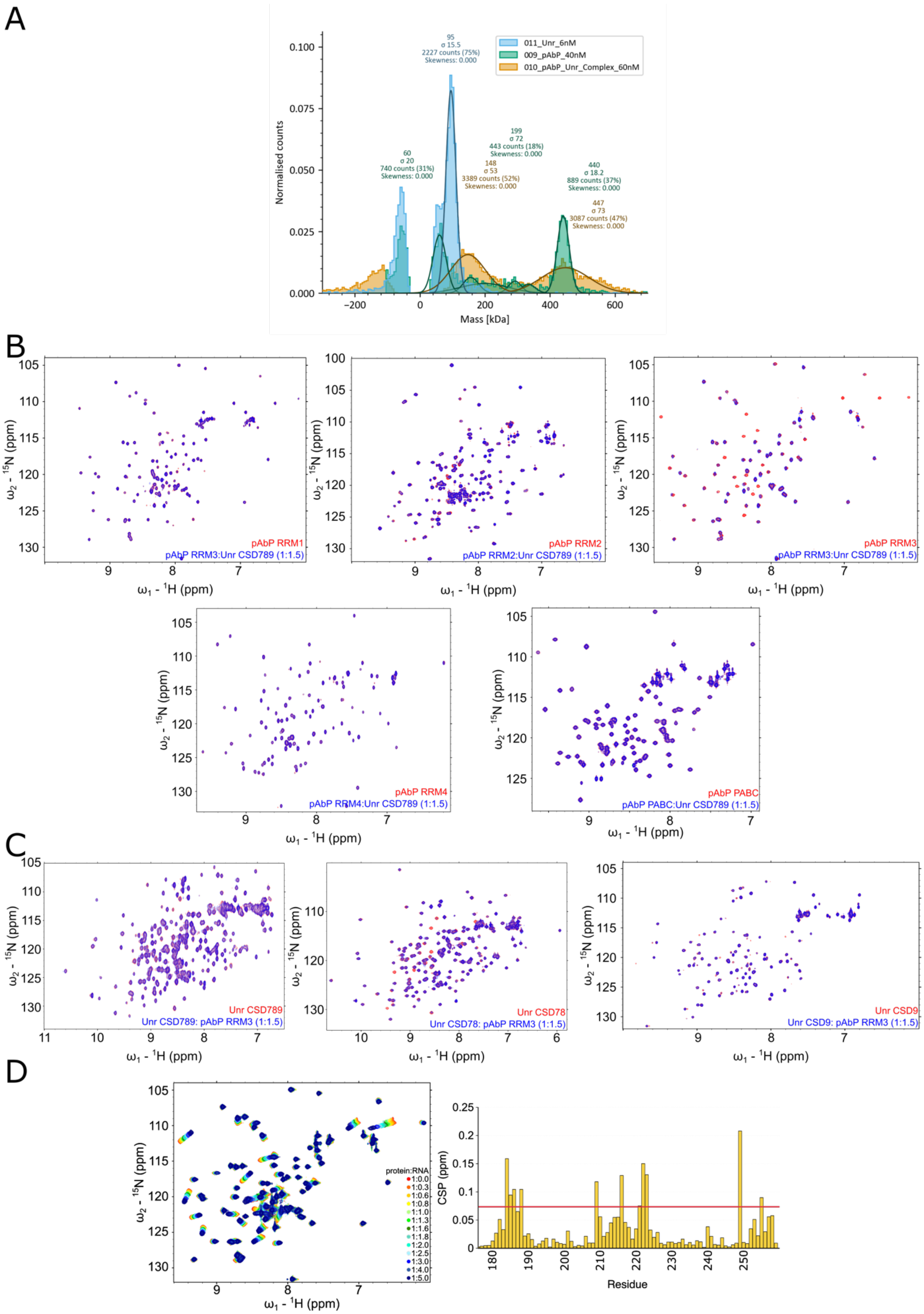
A
0.100
0.075
0.050
0.025
0.000
Normalised counts
Mass [kDa]
-200
0
200
400
600
011_Unr_6nM
009_pAbP_40nM
010_pAbP_Unr_Complex_60nM
95
σ 15.5
2227 counts (75%)
Skewness: 0.000
60
σ 20
740 counts (31%)
Skewness: 0.000
199
σ 72
443 counts (18%)
Skewness: 0.000
148
σ 53
3389 counts (52%)
Skewness: 0.000
440
σ 18.2
889 counts (37%)
Skewness: 0.000
447
σ 73
3087 counts (47%)
Skewness: 0.000
B
ω2 - 15N (ppm)
ω1 - 1H (ppm)
pAbP RRM1
pAbP RRM3:Unr CSD789 (1:1.5)
pAbP RRM2
pAbP RRM2:Unr CSD789 (1:1.5)
pAbP RRM3
pAbP RRM3:Unr CSD789 (1:1.5)
pAbP RRM4
pAbP RRM4:Unr CSD789 (1:1.5)
pAbP PABC
pAbP PABC:Unr CSD789 (1:1.5)
C
Unr CSD789
Unr CSD789: pAbP RRM3 (1:1.5)
Unr CSD78
Unr CSD78: pAbP RRM3 (1:1.5)
Unr CSD9
Unr CSD9: pAbP RRM3 (1:1.5)
D
protein:RNA
1:0.0
1:0.3
1:0.6
1:0.8
1:1.0
1:1.3
1:1.6
1:1.8
1:2.0
1:2.5
1:3.0
1:4.0
1:5.0
CSP (ppm)
Residue
180
190
200
210
220
230
240
250

**Figure S6 a:** Mass photometry histogram showing weight distributions of dUnr and pAbp full length proteins either isolated or as a complex in solution. Only species that have a population size of more than 10% are annotated. **b**: $^{1}H$,$^{15}N$-HSQC spectra show the interaction between different parts of *Drosophila* pAbp (RRM1, RRM2, RRM3, RRM4 and the PABC domain) ($^{15}N$ labeled; red) with 1.5 molar excess of unlabeled CSD789 (blue). **c:** $^{1}H$,$^{15}N$-HSQC spectra show the reverse interaction study of $^{15}N$ labeled CSD789 (left), CDS78 (middle) and CSD9 (right) (all red) with 1.5 molar excess of unlabeled pAbp-RRM3 (red). **d:** $^{1}H$,$^{15}N$-HSQC NMR titration experiments of pAbp-RRM3 and the calculated chemical shift pertubations plot for an A5-mer + at a protein:RNA ration of 1:2. The red line in the CSP plots indicates the average plus the standard deviation of all measured shifts, which was used to identify significant shifts[58].

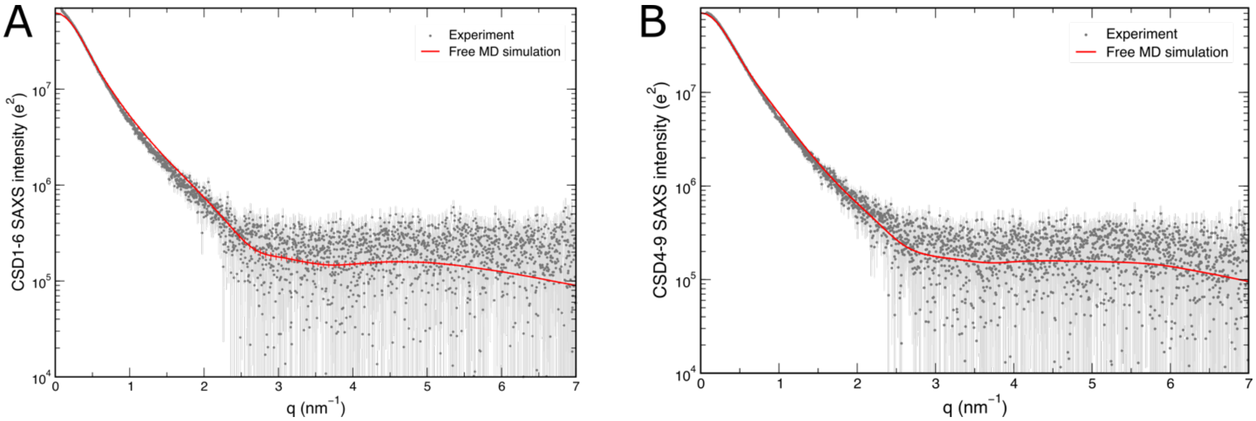


**Figure S7**: The fit of SAXS driven MD simualtions plotted in the scatterig curve of CSD1-6 (a) and CSD4-9 (b) in solution.

**Table S1. Crystal data collection and refinement statistics**

| | **CSD789 + RNA** | **CSD789 + RRM3** |
|---|---|---|
| Wavelength | 0.976 | 0.873 |
| Resolution Range | 48.79 - 1.6 (1.657 - 1.6) | 36.51 - 2.99 (3.1 - 2.99) |
| Space group | C 1 2 1 | P 1 21 1 |
| Unit cell | 97.92 36.78 85.83 90 119.811 90 | 51.6 52.02 73.5 90 96.72 90 |
| Total reflections | 235687 (22904) | 49069 (3278) |
| Unique reflections | 35303 (3434) | 7837 (679) |
| Multiplicity | 6.7 (6.7) | 6.3 (4.8) |
| Completeness (%) | 99.75 (98.76) | 98.08 (84.77) |
| Mean I/sigma(I) | 13.09 (1.31) | 6.56 (2.09) |
| Wilson B-factor | 21.23 | 44.16 |
| R-merge | 0.09 (1.2) | 0.26 (0.66) |
| R-meas | 0.0987 (1.32) | 0.28 (0.74) |
| R-pim | 0.037 (0.50) | 0.11 (0.32) |
| CC 1/2 | 0.99 (0.52) | 0.98 (0.72) |
| CC* | 1 (0.83) | 0.99 (0.92) |
| Reflections used in refinement | 35297 (3434) | 7834 (679) |
| Reflections used for R-free | 1764 (171) | 392 (34) |
| R-work | 0.187 (0.335) | 0.25 (0.315) |
| R-free | 0.210 (0.39) | 0.312 (0.346) |
| CC(work) | 0.960 (0.77) | 0.90 (0.71) |
| CC(free) | 0.95 (0.71) | 0.84 (0.69) |
| Number of non-hydrogen atoms | 2099 | 2296 |
| macromolecules | 1826 | 2279 |
| ligands | 5 | 0 |
| solvent | 268 | 17 |
| Protein residues | 219 | 288 |
| RMS(bonds) | 0.008 | 0.005 |
| RMS(angles) | 1 | 0.76 |
| Ramachandran favored (%) | 98.6 | 94.64 |
| Ramachandran allowed (%) | 0.93 | 4.64 |
| Ramachandran outliers (%) | 0.47 | 0.71 |
| Rotamer outliers (%) | 0 | 4.47 |
| Clashscore | 4.52 | 3.33 |
| Average B-factor | 25.22 | 40.31 |
| macromolecules | 23.6 | 40.35 |
| ligands | 35.42 | |
| solvent | 36.04 | 35.31 |
| Number of TLS groups | | 5 |

Statistics for highest-resolution shell are shown in parentheses

**Table S2. SAXS data collection and refinement statistics**

| | **dCSD789 A15mer** | **dCSD789** | **dCSD1-6** | **dCSD4-9** |
|---|---|---|---|---|
| (a) Sample Details | | | | |
| Organism | *E. coli* BL2 (DE3) | *E. coli* BL2 (DE3) | *E. coli* BL2 (DE3) | *E. coli* BL2 (DE3) |
| Source | *this work* | *this work* | *this work* | *this work* |
| Uniprot sequence ID | Q9VSK3 | Q9VSK3 | Q9VSK3 | Q9VSK3 |
| Description | UNR A759-D990, with TEV-cleaved N-terminal His6-tag in complex with A15mer RNA | UNR A759-D990, with TEV-cleaved N-terminal His6-tag | UNR A176-H677, with TEV-cleaved N-terminal His6-tag | UNR V428-D990, with TEV-cleaved N-terminal His6-tag |
| Molecular mass M from chemical composition (Da) | 30626 | 25750 | 56868 | 63121 |
| loading concentration (mg/ml) | 1.5 and 5.8 mg/ml | 1.6 and 6.2 mg/ml | 0.92 and 3.73 mg/ml | 0.83 and 4.18 mg/ml |
| injection volume (ul) | 30 | 30 | 30 | 30 |
| concentration (uM) | 50/190 | 58/225 | 15/65 | 13/66 |
| Solvent composition and source | 20 mM Hepes/NaOH pH 7.5, 150 mM NaCl and 1 mM DTT | | | |
| (b) SAS data collection parameter | | | | |
| Source and instrument | Hamburg PETRA-III P12 with Dectris Pilatus 6M | Grenoble ESRF BM29 with Dectris Pilatus 1M | | |
| Wavelength (A) | 1.24 | 0.9919 | | |
| Sample-detector distance (m) | 3 | 2,867 | | |
| q-measurement range (nm-1) | 0.0224-7.3176 | 0.0355-4.9391 | 0.0385-4.9335 | |
| Radiation damage monitoring | frame-by-frame comparison | | | |
| Exposure time (s) & number | 0.195x20 | 1.0x10 | | |
| Sample configuration | sample changer with flow through capillary measurement | | | |
| Sample temperature (℃) | 20 | | | |
| (c) Software employed for SAS data reduction, analysis and interpretation | | | | |
| SAXS data processing | I(q) vs. q using Bsx cube, solvent subtraction and curve merging using PRIMUSqt from ATSAS | | | |
| Basic analyses: Guinier, P(r), Vp | PRIMUSqt from ATSAS 2.7.1 | | | |

| | | | | |
|---|---|---|---|---|
| Atomic structure modelling | CRYSOL 2.8.2 from PRIMUSqt in ATSAS 2.8 | EOM 2.1 from PRIMUSqt in ATSAS 2.7.1 | CRYSOL 2.8.2 from PRIMUSqt in ATSAS 2.8 | |
| Molecular graphics | -- | PyMol v3.3.1 (Linux) | -- | -- |
| | | (d) Structural parameters | | |
| | | Guinier analysis | | |
| I(0) (raw) | 29+/-0.05 | 26.74+/-0.04 | 56.04+/-0.17 | 58.27+/-0.19 |
| $R_g$ (Å) | 22.3+/-0.02 | 23.4+/-0.07 | 43.6+/-0.31 | 49+/-0.34 |
| $qR_g$ max ($q_{min}$ = 0.0066 Å$^{-1}$) | 1.3 | 1.29 | 1.25 | 1.13 |
| Coefficient of correlation, $R^2$ | 0.96 | 0.98 | 0.86 | 0.94 |
| | | P(r) Analysis from AUTOGNOM | | |
| I(0) (cm-1) | 28.9 | 26.51 | 55.19 | 48.99 |
| $R_g$ (Å) | 22.6 | 23.1 | 43.3 | 38.1 |
| $d_{max}$ (Å) | 75.2 | 70.7 | 141.1 | 107.2 |
| $q$ range (Å$^{-1}$) | 0.107-3.585 | 0.092-3.423 | 0.147-1.833 | 0.0714-1.631 |
| $\chi^2$ (total estimate from *GNOM*) | 0.88 | 0.71 | 0.6 | 0.59 |
| Porod volume (Å$^{-3}$) (ratio $V_P$/calculated *M)* | 36490 | 37470 | 148820 | 132810 |
| | | (f) Atomistic modelling | | |
| Method | | EOM | CRYSOL | |
| Crystal stucture | | UNR CSD78 and 9 (PDB ID: 6y69, 6y4h) | | |
| Flexible linker definition | | V919-R920 | | |
| *EOM* | | default parameters, 10 000 models in initial ensemble, native-like models, constant subtraction allowed | | |
| $\chi^2$ | | 0.932 | | |
| Constant subtraction | | 0.018 | | |
| No. of representative structures | | 3 | | |

**Table S3. Distance restraints used for SAXS-driven MD simulation**

**Unr CSD789**

| restr. ID | Res1 | Atom(s) | Res2 | Atom(s) |
|---|---|---|---|---|
| 1 | ARG-765 | HB1, HB2 | ILE-837 | HD1, HD1, HD3 |
| 2 | ARG-765 | HD1, HD2 | ILE-837 | HD1, HD2, HD3 |

| | | | | |
|---|---|---|---|---|
| 3 | PHE-767 | HE1, HE2 | ILE-837 | HB |
| 4 | PHE-767 | HD1 | ILE-837 | HB |
| 5 | PHE-767 | HE1, HE2 | ILE-837 | HD1, HD2, HD3 |
| 6 | PHE-767 | HD1 | ILE-837 | HD1, HD2, HD3 |
| 7 | PHE-767 | HE1, HE2 | ILE-837 | HG11, HG12 |
| 8 | PHE-767 | HD1 | ILE-837 | HG11, HG12 |
| 9 | PHE-767 | HE1, HE2 | ILE-837 | HG21, HG22, HG23 |
| 10 | PHE-767 | HD1 | ILE-837 | HG21, HG22, HG23 |
| 11 | LEU-781 | HD11, HD12, HD13 | LEU-781 | HA |
| 12 | LEU-781 | HD11, HD12, HD13 | ILE-837 | HD1, HD2, HD3 |
| 13 | LEU-781 | HB1, HB2 | ILE-837 | HD1, HD2, HD3 |
| 14 | GLU-806 | HG1, HG2 | ILE-837 | HD1, HD2, HD3 |
| 15 | GLU-806 | HB1, HB2 | ILE-837 | HD1, HD2, HD3 |
| 16 | ILE-837 | HB | THR-888 | HG21, HG22, HG23 |
| 17 | ILE-837 | HG11, HG12 | THR-888 | HG21, HG22, HG23 |
| 18 | TYR-865 | HE1, HE2 | ALA-769 | HB1, HB2, HB3 |
| 19 | ILE-887 | HG21, HG22, HG23 | GLY-804 | HA1 |
| 20 | ILE-887 | HG21, HG22, HG23 | ALA-769 | HA |
| 21 | ILE-887 | HB | ALA-769 | HA |
| 22 | ILE-887 | HD1, HD2, HD3 | ALA-769 | HA |
| 23 | ILE-887 | HG21, HG22, HG23 | PHE-767 | HB1, HB2 |
| 24 | ILE-887 | HG21, HG22, HG23 | ALA-769 | HB1, HB2, HB3 |
| 25 | ILE-887 | HB | ALA-769 | HB1, HB2, HB3 |
| 26 | ILE-887 | HD1, HD2, HD3 | ALA-769 | HB1, HB2, HB3 |
| 27 | ILE-887 | HG21, HG22, HG23 | GLU-779 | HB1, HB2, HB3 |
| 28 | ILE-887 | HB | PHE-767 | HD1 |
| 29 | THR-888 | HG21, HG22, HG23 | PHE-767 | HB1, HB2 |
| 30 | THR-888 | HG21, HG22, HG23 | PHE-767 | HD1 |
| 31 | THR-888 | HG21, HG22, HG23 | PHE-767 | HE1, HE2 |
| 32 | PHE-767 | HD1 | ILE-887 | HB |
| 33 | PHE-767 | HD1 | ILE-887 | HG11, HG12 |
| 34 | PHE-767 | HD1 | ILE-887 | HG21, HG22, HG23 |
| 35 | PHE-767 | HB1, HB2 | ILE-887 | HG21, HG22, HG23 |
| 36 | PHE-767 | HB1, HB2 | THR-888 | HG21, HG22, HG23 |
| 37 | PHE-767 | HE1, HE2 | THR-888 | HG21, HG22, HG23 |
| 38 | PHE-767 | HD1 | THR-888 | HG21, HG22, HG23 |
| 39 | PHE-767 | HZ | THR-888 | HG21, HG22, HG23 |
| 40 | ALA-769 | HB1, HB2, HB3 | ILE-887 | HB |
| 41 | ALA-769 | HA | ILE-887 | HD1, HD2, HD3 |
| 42 | ALA-769 | HB1, HB2, HB3 | ILE-887 | HD1, HD2, HD3 |

| | | | | |
|---|---|---|---|---|
| 43 | ALA-769 | HB1, HB2, HB3 | TYR-865 | HE1, HE2 |
| 44 | ALA-769 | HB1, HB2, HB3 | ILE-887 | HG21, HG22, HG23 |
| 45 | ALA-769 | HA | ILE-887 | HG21, HG22, HG23 |
| 46 | GLU-779 | HB1, HB2 | ILE-887 | HG21, HG22, HG23 |
| 47 | LEU-803 | HD21, HD22, HD23 | ILE-887 | HD1, HD2, HD3 |
| 48 | LEU-803 | HD21, HD22, HD23 | ILE-887 | HD1, HD2, HD3 |
| 49 | GLY-804 | HA1, HA2 | ILE-887 | HG11 |
| 50 | GLY-804 | HA1, HA2 | ILE-887 | HG12 |
| 51 | GLY-804 | HA1, HA2 | ILE-887 | HG21, HG22, HG23 |
| 52 | LEU-781 | HD21, HD22, HD23 | ILE-837 | HA |
| 53 | LEU-781 | HD21, HD22, HD23 | ILE-837 | HD1, HD2, HD3 |

**UNR CSD789 + RNA**

| restr. ID | Res1 | Atom(s) | Res2 | Atom(s) |
|---|---|---|---|---|
| 1 | GLN-898 | OE1 | GLN-975 | HN |
| 2 | GLN-898 | OE1 | GLN-975 | NE2 |
| 3 | GLN-898 | CB | GLN-975 | NE2 |
| 4 | GLN-898 | OE1 | ADE-9 | CG |
| 5 | LYS-979 | HD1 | ADE-9 | C6 |
| 6 | LYS-979 | HZ3 | ADE-9 | N3 |
| 7 | LYS-979 | HZ1 | ADE-9 | O2' |
| 8 | ARG-976 | HH21 | ASP-861 | HB1 |
| 9 | ASN-977 | HD21 | ADE-6 | N6 |
| 10 | ASN-977 | HD21 | ADE-6 | N1 |
| 11 | ARG-856 | HE | ADE-5 | H62 |
| 12 | ARG-856 | CG | ADE-5 | N6 |
| 13 | GLU-786 | CG | ADE-5 | H52 |
| 14 | GLU-786 | OE1 | ADE-5 | H62 |
| 15 | GLU-786 | OE1 | ARG-856 | HE |
| 16 | GLU-786 | OE2 | ARG-856 | NH2 |
| 17 | PRO-860 | CG | ADE-5 | C4 |
| 18 | PRO-860 | HB1 | ADE-5 | C5 |
| 19 | PRO-860 | CA | ADE-5 | C6 |
| 20 | PRO-860 | O | ADE-5 | N6 |

### Table S4. ITC data parameters

| Name | [Syr] (M) | [Cell] (M) | N (sites) | KD (M) | ΔH (kcal/mol) | ΔG (kcal/mol) | -TΔS (kcal/mol) | Number of replicates | comments |
|---|---|---|---|---|---|---|---|---|---|
| CSD789 (cell) versus poly(A)-7mer (syringe) | $510 \cdot 10^{-6}$ | $60 \cdot 10^{-6}$ | 0.47 ± 0.16 | $17.7 \pm 5.2 \cdot 10^{-6}$ | -17.1 ± 3.6 | -6.4 ± 0.2 | 10.8 ± 3.7 | 3 | |
| CSD789 (cell) versus poly(A)-8mer (syringe) | $200 \cdot 10^{-6}$ | $20 \cdot 10^{-6}$ | 0.61 ± 0.24 | $4.4 \pm 2.7 \cdot 10^{-6}$ | -12.4 ± 2.7 | -7.2 ± 0.4 | 5.1 ± 2.8 | 4 | |
| CSD789 (cell) versus poly(A)-9mer (syringe) | $300 \cdot 10^{-6}$ | $60 \cdot 10^{-6}$ | 0.58 ± 0.36 | $4.3 \pm 1.3 \cdot 10^{-6}$ | -14.5 ± 4.7 | -7.2 ± 0.2 | 7.3 ± 4.8 | 4 | |
| CSD789 (cell) versus poly(A)-15mer (syringe) | $200 \cdot 10^{-6}$ | $60 \cdot 10^{-6}$ | 0.31 ± 0.01 | $4.8 \pm 0.8 \cdot 10^{-6}$ | -28.5 ± 12.7 | -7.1 ± 1.0 | 21.4 ± 12.8 | 3 | slight precipitation |
